# Increased vowel contrast and intelligibility in connected speech induced by sensorimotor adaptation

**DOI:** 10.1101/2024.08.04.606537

**Authors:** Sara D. Beach, Sophie A. Johnson, Benjamin Parrell, Caroline A. Niziolek

## Abstract

Alterations to sensory feedback can drive robust adaptive changes to the production of consonants and vowels, but these changes often have no behavioral relevance or benefit to communication (e.g., making “head” more like “had”). This work aims to align the outcomes of adaptation with changes known to increase speech intelligibility – specifically, adaptations that increase the acoustic contrast between vowels in running speech. To this end, we implemented a vowel centralization feedback perturbation paradigm that pushes all vowels towards the center of vowel space, making them sound less distinct from one another. Speakers across the adult lifespan adapted to the centralization perturbation during sentence production, increasing the global acoustic contrast among vowels and the articulatory excursions for individual vowels. These changes persisted after the perturbation was removed, including after a silent delay, and showed robust transfer to words that were not present in the sentences. Control analyses demonstrated that these effects were unlikely to be due to explicit pronunciation strategies and occurred in the face of increasingly more rapid and less distinct production of familiar sentences. Finally, sentence transcription by crowd-sourced listeners showed that speakers’ vowel contrast predicted their baseline intelligibility and that experimentally-induced increases in contrast predicted intelligibility gains. These findings establish the validity of a sensorimotor adaptation paradigm to implicitly increase vowel contrast and intelligibility in connected speech, an outcome that has the potential to enhance rehabilitation in individuals who present with a reduced vowel space due to motor speech disorders, such as the hypokinetic dysarthria associated with Parkinson’s disease.

**Significance statement:** During speech production, humans learn from perceived errors by adjusting their speech to compensate, a process known as sensorimotor adaptation. Here we leveraged sensorimotor adaptation to cause an increase in the distinctiveness of speech sounds. Specifically, speakers rapidly and implicitly learned vowel-specific articulation changes during sentence production, with positive effects on their intelligibility. This study is important for models of sensorimotor integration across all motor domains, as it demonstrates the simultaneous learning of multiple opposing transformations in the course of continuous movement. Our findings show that sensorimotor adaptation can be successfully applied to drive changes relevant to a vital ecological behavior: intelligible communication.

## Introduction

Speech production, like other motor behaviors, responds to changes in sensory feedback. Real-time, external perturbations of auditory feedback, such as raising or lowering the resonant frequencies of the vocal tract (formants), can drive individuals to alter their pronunciation to oppose the perturbation and correct for the error (Houde & Jordan, 1998; Purcell & Munhall, 2006; Villacorta et al., 2007; reviewed in Caudrelier & Rochet-Capellan, 2019). Such *sensorimotor adaptation* has historically been studied with uniform perturbations applied to relatively simple movements in separate trials, whether for speech (isolated words), upper limb control (center-out reaches), or eye movement (saccades). However, natural speech is complex, dynamic, and combinatorial, and causing “head” to sound more like “had,” for example, has little relevance to functional communication.

The acoustic contrast among vowels underlies their perceptual distinctiveness, and hence is key to speech intelligibility (Bradlow et al., 1996; Neel, 2008; Ferguson & Kewley-Port, 2007). Correspondingly, many motor disorders that impair intelligibility, such as Parkinson’s disease, are characterized by reduced vowel contrast (Skodda et al., 2011; Sapir et al., 2010; Lee et al., 2017; Weismer et al., 2001). Rehabilitating this deficit requires motor learning of new speech behaviors through practice or experience (Maas et al., 2008). Because sensorimotor adaptation to altered feedback is both rapid and largely unconscious (Hantzsch et al., 2022; Munhall et al., 2009), a perturbation that could drive speakers to produce more distinct vowels has the potential to increase speech intelligibility without the need for extended practice and the use of explicit cognitive strategies.

Our recent work has shown that speakers do indeed adapt to increase their vowel contrast in response to a nonuniform feedback perturbation during the production of isolated monosyllabic words: as the perturbation pushes all vowels towards the center of formant space (**Figure 1A**), individuals respond by producing corner vowels /i/, /æ/, /ɑ/, and /u/ farther from that center, increasing global measures of vowel contrast (Parrell & Niziolek, 2021). However, most communication occurs in connected speech, and it is not yet clear whether such nonuniform adaptations can be learned as formant trajectories continuously traverse the full acoustic workspace. Humans are capable of learning complex, co-occurring, and indeed opposing motor adaptations (Rochet-Capellan & Ostry, 2011; Parrell & Niziolek, 2021; Zeng et al., 2023), as well as a uniform formant shift applied across the entire vowel space of connected speech (Lametti et al., 2018), but these two paradigms – nonuniform perturbation and continuous movement – have yet to be tested together in speech or, indeed, in any motor domain.

**Figure 1.**
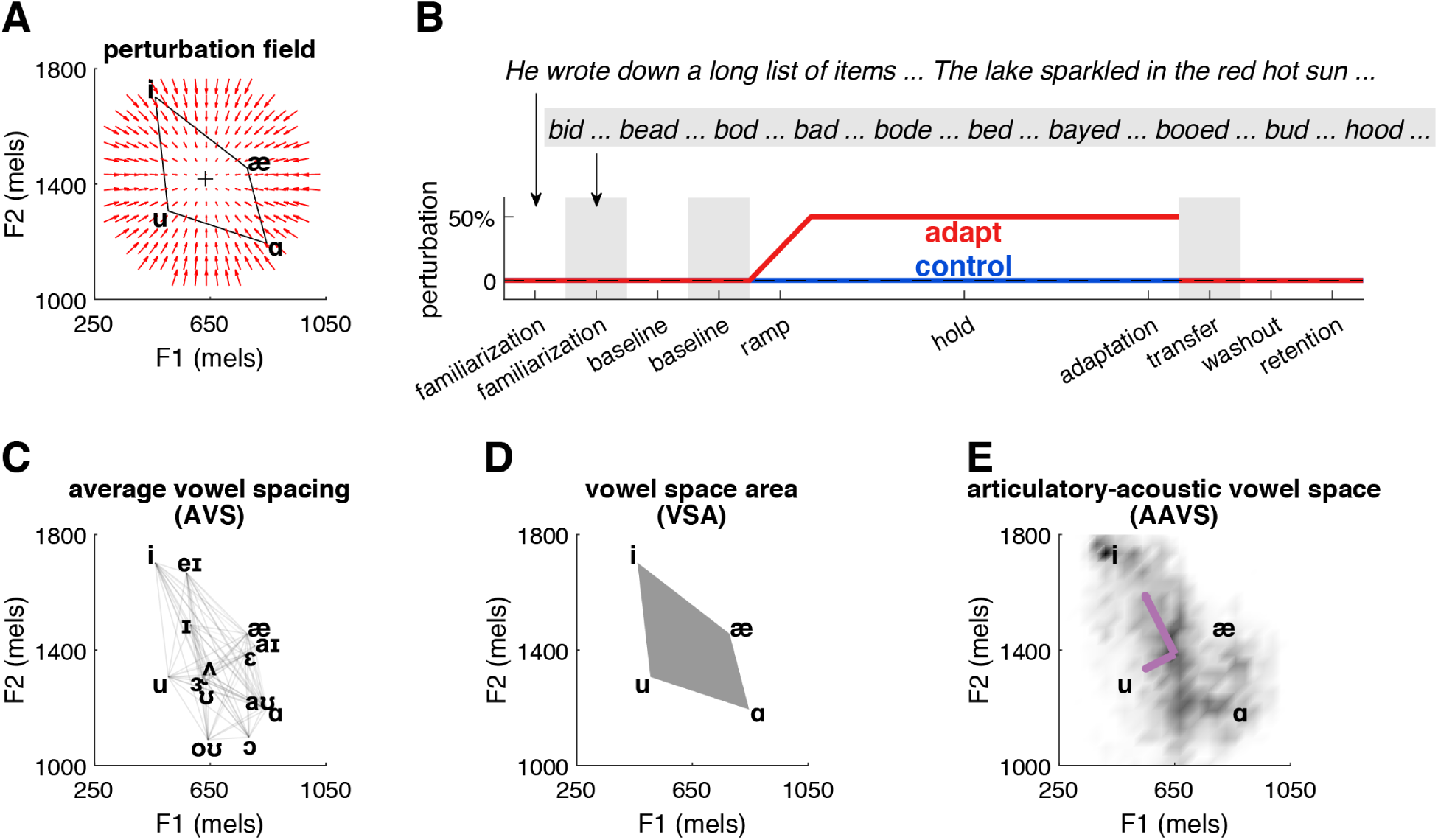
Experimental design and vowel-space outcome metrics. **(A)** Illustration of the perturbation field applied to speech, in which vowels were pushed toward the center of the speaker’s vowel space, indicated by the “+”. This acoustic space is defined by the first two formants, or resonances, of the vocal tract (F1-F2 space). **(B)** Phases of the experiment in adapt and control sessions; the structure across sessions was identical but the perturbation (50% of the Euclidean distance between current F1 and F2 values and the center) was applied only in the adapt session. At top, examples of sentence stimuli. During the shaded phases, stimuli were isolated transfer words representing 10 different English vowels. **(C)** Average vowel spacing (AVS) is the average of all pairwise vowel-to-vowel distances in F1-F2 space. Shown are the 14 vowels of interest in the sentence stimuli. **(D)** Vowel space area (VSA) is the area of the quadrilateral formed by the four corner vowels in F1-F2 space (gray polygon). **(E)** Articulatory-acoustic vowel space (AAVS) is the square root of the generalized variance of all sampled F1 and F2 values (grayscale density), representing formant variability. Purple arrows indicate the eigenvectors of the corresponding covariance matrix. Vowel data in panels A and C–E are from a single representative participant.

Here, healthy speakers across the adult lifespan (ages 18 to 76) read aloud a set of 40 phonetically balanced sentences covering the vowel space of American English (IEEE, 1969). After a baseline phase with normal feedback, vowel centralization was gradually introduced and then held at maximum magnitude for six blocks, the final block of which served as the phase from which adaptation was measured (**Figure 1B**). Auditory feedback was returned to normal in a subsequent washout phase and in a retention phase following a ten-minute silent delay. Additionally, noise-masked transfer phases before the ramp and after the hold phase (shaded regions in **Figure 1B**) assessed changes in the production of isolated words that did not appear in the sentences (Lametti et al., 2018). To control for potential speech changes over the course of the experiment, speakers also completed a control session, identical in structure to the adapt session and counterbalanced in order, in which no auditory perturbations were applied. The primary acoustic index of vowel contrast was *average vowel spacing* (AVS), the average, in mels, of all pairwise vowel-to-vowel Euclidean distances in formant space (**Figure 1C**), a metric that is both sensitive to adaptation effects (Parrell & Niziolek, 2021) and a prevalent clinical measure (Ng & Woo, 2021; Lane et al., 2001). To evaluate potential changes in intelligibility, we presented a mix of baseline and adapted speech recordings – degraded with noise to lower baseline intelligibility – for transcription and word identification by crowd-sourced listeners.

To preview our results, we establish the capacity of sensorimotor adaptation to increase vowel contrast during connected, sentence-level speech, with positive effects on speaker intelligibility. Moreover, this learning persists after the perturbation is removed, transfers to untrained words, occurs across the lifespan, is reflected in both global and vowel-specific changes, and is unlikely to be due to explicit, effortful pronunciation strategies. These findings demonstrate that sensory predictions are highly specific to parts of the workspace traversed during continuous movements, and that those continuous movements can be adapted to complex, nonuniform perturbations during high-level ecological behavior.

## Results

### Speakers adapt to centralization by increasing vowel contrast

Speakers responded to the centralization perturbation by increasing AVS in the adapt session but not the control session (main effect of *session type*: (*F*(1,245) = 7.96, *p* = 0.01; **Figure 2A**). There was no effect of *phase* (*F*(2,245) = 0.46, *p* = 0.63) and a nonsignificant *session type* × *phase* interaction (*F*(2,245) = 3.04, *p* = 0.0537), indicating that AVS stayed elevated from the adaptation phase through the washout phase (when the perturbation was removed) and the retention phase (after a ten-minute silent wait) (see also **Figure 2B**). Overall, across these three phases, AVS was 2.17±6.89% (M±SD) above baseline in the adapt session, in comparison to −0.65±5.02% (below baseline) in the control session. However, after correction for multiple comparisons, none of the six phases across adapt and control was significantly different from baseline (*p*’s ≥ 0.03).

**Figure 2.**
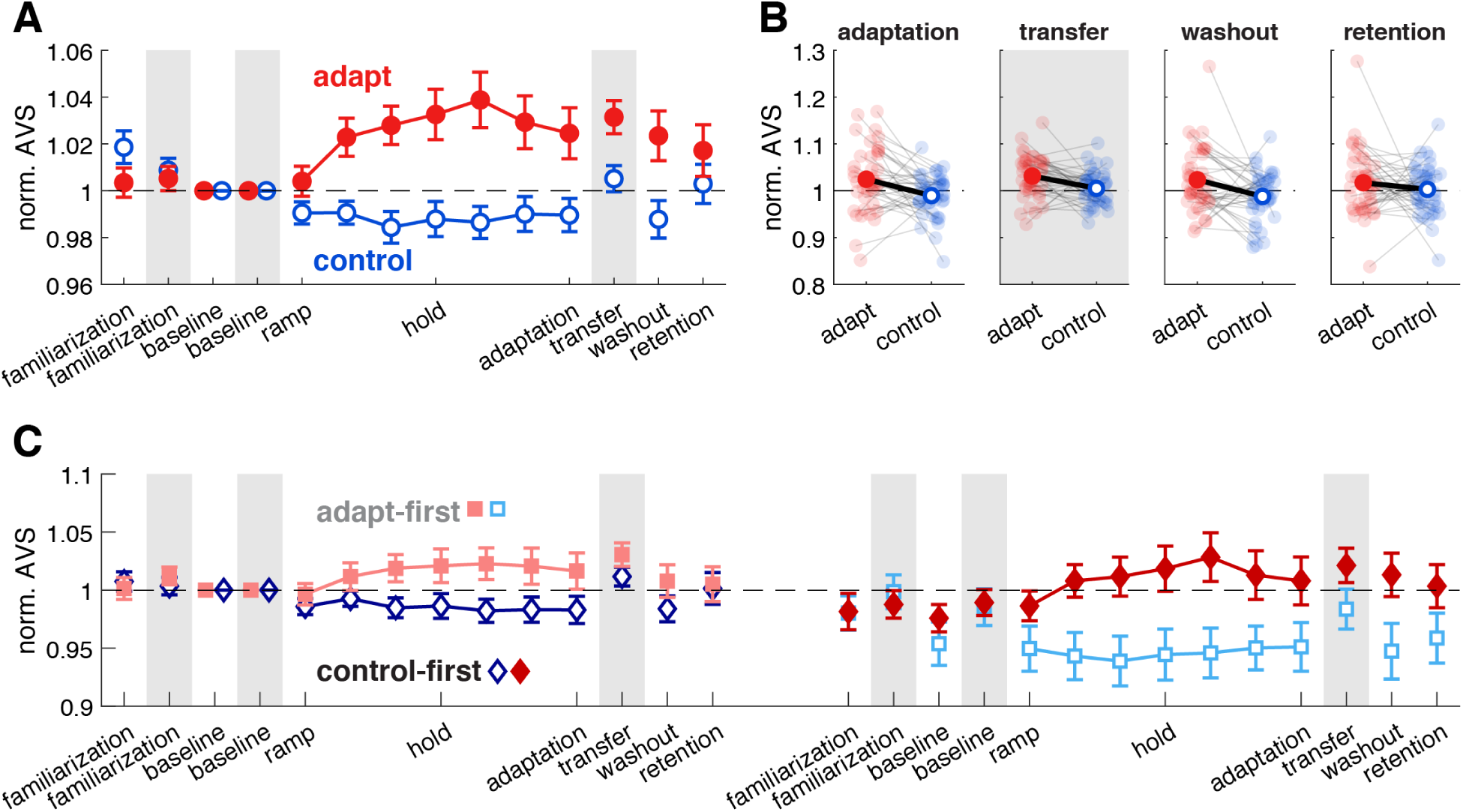
Adaptation increases global vowel contrast. **(A)** AVS collapsed across first and second sessions and normalized to the within-session baseline shows that vowel contrast increases in adapt (red) but not control (blue) sessions, and that this adaptation generalizes to untrained transfer words (shaded regions). **(B)** Individual data (small markers connected by gray lines) and group means for within-session normalized AVS in each phase of interest. **(C)** AVS for first and second sessions separately, normalized to the first-session baseline, reveals wider differences between adapt and control AVS in the second session. Data from adapt-first participants are pink squares followed by light blue squares. Data from control-first participants are dark blue diamonds followed by red diamonds. Error bars show standard error.

The lack of a *session type* × *session order* interaction (*F*(1,245) = 2.58, *p* = 0.12) suggested that sensorimotor adaptation drove an increase in vowel contrast no matter whether participants experienced centralization in their first or second session (due to counterbalancing). However, we performed an exploratory analysis to investigate potential effects of returning to the lab for a second session to produce familiar sentences. We normalized individuals’ vowel-space measures to their baseline value in the *first* session and performed the same ANOVA to test for differences between first and second sessions.

The significant effect of *session type* (main effect: *F*(1,245) = 13.18, *p* < 0.001) had a nonsignificant interaction with *session order* (interaction: *F*(1,245) = 3.39, *p* = 0.07); specifically, the differences between adapt and control AVS were numerically but not significantly greater in the second session (**Figure 2C**). This was primarily driven by AVS contraction in the second control session (−4.77±9.77%), suggesting that, in the absence of the perturbation, familiarity drives speakers to a casual manner of speaking characterized by reduced vowel contrast. Post-hoc tests on the *session type* × *phase* interaction (*F*(2,245) = 3.19, *p* = 0.0466) revealed differences between adapt and control AVS in each phase of interest: adaptation (*F*(1,81) = 15.39, *p* < 0.001), washout (*F*(1,81) = 13.50, *p* < 0.001), and retention (*F*(1,81) = 4.24, *p* = 0.0463), and there was no three-way interaction with *session order* (*F*(2,245) = 0.26, *p* = 0.77). In sum, with respect to speakers’ first-session baseline AVS, exposure to the centralization perturbation prevented a natural contraction of the vowel space over time and repetitions.

To permit comparison with the wider speech literature, we quantified adaptation with two additional vowel-space metrics, vowel space area (VSA) and articulatory-acoustic vowel space (AAVS). VSA (**Figure 1D**) is the area of the quadrilateral formed by the four corner vowels. Although VSA revealed no significant effects or interactions with either normalization procedure at our predetermined measurement point, a numerical increase under adaptation was evident and reached significance (above the within-session baseline) in blocks one and three of the hold phase (*p*’s < 0.04; **Figure S1**). AAVS (**Figure 1E**) is a trajectory-based measure reflecting the formant distribution in F1-F2 space (Whitfield & Goberman, 2014). For AAVS using within-session normalization, the effect of *session type* (main effect: *F*(1,245) = 4.60, *p* = 0.04) interacted with *session order* (interaction: *F*(1,245) = 6.73, *p* = 0.01; **Figure S2**). Follow-up ANOVAs revealed no significant effects or interactions in the first session, but a main effect of *session type* in the second session (*F*(1,122) = 12.07, *p* < 0.001). Specifically, the second-session AAVS revealed a dramatic and persistent increase in working articulatory space under adaptation: 10.66±17.79% above baseline in the adapt session, in comparison to 1.67±8.79% in the control session. Consistent with no effect of or interaction with *phase* (*p*’s > 0.86), AAVS expansion in the adaptation phase (10.17%) was maintained during the washout (11.65%) and retention phases (10.18%). After correction for six comparisons, only the adaptation phase of the adapt session significantly differed from baseline (*t*(20) = 3.14, *p* = 0.01, *d* = 0.42). With the first-session normalization, AAVS showed a main effect of *session type* (*F*(1,245) = 11.51, *p* < 0.01) and a nonsignificant interaction with *phase* (*F*(2,245) = 2.98, *p =* 0.0568), confirming that adaptation increased articulatory-acoustic working space.

### Increased vowel contrast transfers to untrained words

The increase in AVS showed a robust transfer to isolated words that did not appear in the sentence stimuli and were never produced with the perturbation (shaded regions in **Figure 2**), as indicated by a main effect of *session typ*e (*F*(1,81) = 13.04, *p* < 0.001). AVS in the transfer phase was significantly different from its baseline in the adapt session (3.14±4.52%; *t*(40) = 4.45, *p* < 0.001, *d* = 0.68) but not the control session (0.05±3.57%; *t*(40) = 0.92, *p* = 0.36, *d* = 0.14), correcting for two comparisons.

The VSA metric also revealed significant transfer (main effect of *session type*: *F*(1,81) = 6.69, *p* = 0.01): VSA was 7.38±13.47% above baseline in the adapt session, in comparison to 1.41±11.97% in the control session (**Figure S1**). (AAVS, a measure based on formant trajectories in connected speech, was not calculated for transfer words produced in isolation.) An exploratory first-session normalization confirmed these results, with significant main effects of *session type* (AVS: *F*(1,81) = 7.59, *p* = 0.01; VSA: *F*(1,81) = 4.17, *p* = 0.0479) and no effects of or interactions with *session order* (*p*’s > 0.07). Overall, each metric revealed a generalization of expanded vowel spacing in connected speech to untrained single words.

### Smaller vowel spaces undergo greater adaptation

With an eye towards future clinical utility, we investigated the relationships among speaker demographics, baseline vowel space, and sensorimotor adaptation using stepwise linear regression. At baseline of their first session, younger participants (*p* = 0.04) and males (*p* < 0.01) presented with smaller vowel spaces (raw sentence-based AVS in mels; *F*(2,38) = 9.59, *p* < 0.001, R^2^ = 0.34). There was no interaction between *age* and *sex* (*p* = 0.25).

When we entered *age*, *sex*, and *baseline AVS* into a second model, only *baseline AVS* was a significant predictor of future vowel-space gains under adaptation (*F*(1,39) = 5.67, *p* = 0.02, R^2^ = 0.13). Specifically, individuals who presented with smaller vowel spaces showed larger increases in AVS in the adaptation phase of their adapt session, up to a maximum of 117% of their baseline value. The model estimated that an individual whose vowels were, on average, 10 mels closer together could expect 0.90% more AVS expansion. These regression results held when gain was calculated as a raw increase in mels: *baseline AVS* was its only significant predictor (*F*(1,39) = 4.78, *p* = 0.03, R^2^ = 0.11). The model estimated that an individual whose vowels were, on average, 10 mels closer together could expect an additional 2.3 mels of AVS gain. These results suggest that vowel contrast can increase even in healthy speakers with typical baseline vowel spaces, although speakers who habitually use a smaller working space may have more room for expansion.

### Speakers learn multiple vowel-specific adaptations

In order to counteract the centralization perturbation and increase vowel contrast, speakers must simultaneously learn multiple vowel-specific compensatory adaptations in the course of running speech. To quantify vowel-specific formant change (distinct from the global AVS measure) for each of 14 vowels of interest in the sentence productions, we calculated its distance from the center of the participant’s vowel space at baseline (gray lines in **Figure 3A**; a representative participant) and in the adaptation phase (colored lines). We evaluated whether this distance increased over time (i.e., from baseline to the adaptation phase) in the adapt relative to the control session, as would be expected of change opposing the perturbation.

**Figure 3.**
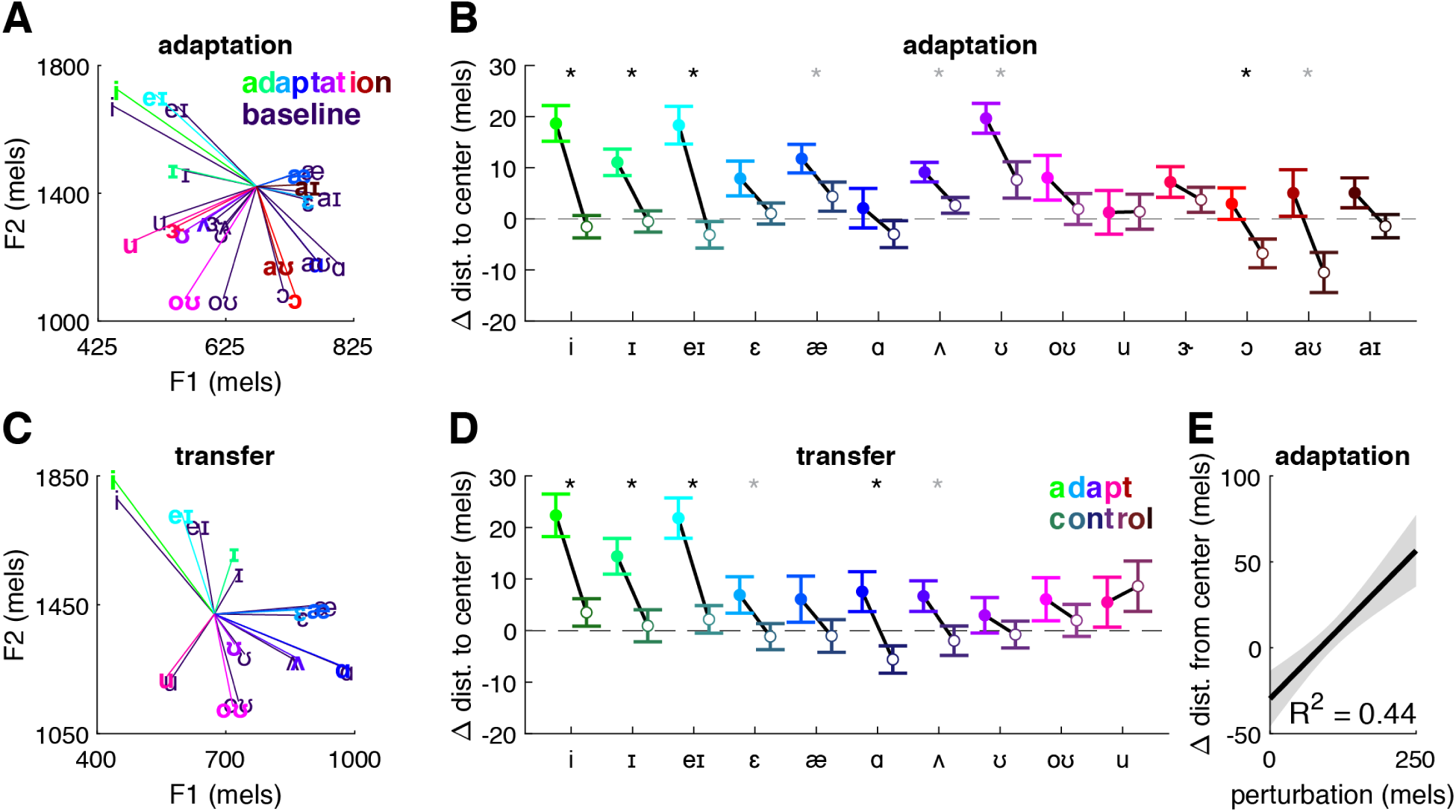
Vowel-specific adaptation. **(A)** Illustration of individual vowels’ increased distance from the center of the vowel space from baseline (gray) to the adaptation phase (sentence production) of the adapt session (colors). Data are from the same representative participant as in Figure 1. **(B)** Group means (±SE) for each sentence vowel in the adaptation phase of adapt (bright) and control (dark) sessions show the change from baseline in the distance to the center of vowel space. Gray * = significant difference between adapt and control sessions, paired *t*-test. Black * = significant after Holm-Bonferroni correction. **(C)** As in (A), vowels from the baseline and transfer phases of the adapt session. Data are from the same representative participant. **(D)** As in (B), vowels from the transfer phase of the adapt session. **(E)** Fitted model and 95% confidence interval for vowels’ change in distance from the center of the vowel space as a function of the average perturbation size applied to that vowel.

**Figure 3B** shows that only a subset of the 14 sentence vowels were produced farther from the center of the vowel space in the adapt (bright colors) vs. the control session (dark colors). The significant effect of *session type* (main effect: *F*(1,1147) = 19.85, *p* < 0.001) varied by *vowel* (interaction: *F*(13,1147) = 2.76, *p* < 0.001); there was also a main effect of *vowel* (*F*(13,1147) = 4.92, *p* < 0.001). Post-hoc comparisons revealed that these changes in distance-to-center over time differed between adapt and control sessions in eight of the vowels (paired *t*-tests, *p*’s < 0.05); four remained significant after correction for multiple comparisons: /i/, /ɪ/, /e/, and /ɔ/. Furthermore, the *direction* of the change in distance-to-center was in line with our hypothesis: across all vowels measured in adapt and control sessions, 11 of them showed non-zero change from baseline (*t*-tests against 0, *p*’s < 0.05); once corrected for 28 comparisons, six remained significant: /i/, /ɪ/, /e/, /æ/, /ʌ/, and /ʊ/. In all cases, these were adapt-session vowels that moved away from the center of the vowel space (i.e., positive values in Figure 3B), thereby opposing the perturbation. Incidentally, the adapt-vs.-control differences identified in /ɔ/ and /aʊ/ were driven by movement towards the center in the control session, suggesting that, in these vowels, adaptation counters a natural drift towards less distinct pronunciations over time and repetitions.

Vowel-specific adaptation was confirmed by testing the ten vowels in the transfer words in the same way (Figure 3C; same representative participant). The effect of *session type* (*F*(1,819) = 17.98, *p* < 0.001) also varied by *vowel* (interaction: *F*(9,819) = 2.73, *p* < 0.01); there was additionally a main effect of *vowel* (*F*(9,819) = 3.68, *p* < 0.001). Post-hoc comparisons revealed that changes in distance-to-center over time differed between adapt and control sessions in six vowels; four remained significant after correction (Figure 3D). The direction of the effect matched that of the sentence-vowel results described above: five vowels showed non-zero change from baseline; after correction for 20 comparisons, three adapt-session vowels showed significant movement away from the center of the vowel space (/i/, /ɪ/, and /e/). The adapt-vs.-control difference in /ɑ/ was driven by movement towards the center in the control session.

Because vowels at the edges of vowel space received larger perturbations (50% of their distance to center), we tested whether perturbation magnitude could explain the observed vowel-specific adaptation effects. Indeed, linear mixed effects modeling of the sentence vowels revealed that larger perturbations were positively associated with greater adaptation (*F*(1,17.94) = 30.20, *p* < 0.001), with a model R^2^ of 0.44 (Figure 3E).

In sum, the fact that different vowels demonstrated adaptation by moving in nonuniform directions in vowel space indicates that speakers can learn multiple compensatory changes in the course of connected speech.

### Increased vowel contrast under adaptation is not merely clear speech

Changes in vowel production in both adapt and control sessions, measured both individually and globally with AVS, suggested that simply repeating the same utterances led to reduced vowel contrast. Acoustic correlates of such “repetition reduction”, in addition to reduced vowel spacing, include shorter durations (Jacobs et al., 2015), reduced intensity (Lam & Watson, 2010), and flatter pitch (Baker & Bradlow, 2009), all serving to make the repeated utterance less prominent. On the other hand, speakers can and do *enhance* these features when aiming to speak more clearly, for example, when speaking to an individual with hearing difficulties or in a noisy environment, or when conveying information that a listener cannot predict from context (Baker & Bradlow, 2009; Scarborough, 2010; Oviatt et al., 1998; Burnham et al., 2010; Lindblom, 1990). Importantly, “hyperarticulation” in clear speech is marked not only by increases in vowel space, but also by changes in these other aspects of speech, and particularly by increases in duration, which allow more time for articulatory movement (Lindblom, 1990).

To rule out the possibility that observed changes in vowel spacing (Figures 2 and **3**) were due to speakers’ attempts to speak clearly rather than to sensorimotor adaptation, we analyzed the duration, peak intensity, and vocal pitch (both maximum and range) of utterances from the baseline to the adaptation phase. If pronunciation changes were due to hyperarticulation, we expected to see duration (and potentially intensity and pitch: Burnham et al., 2010; Krause & Braida, 2004) increase in the adapt session relative to the control session.

In fact, sentence duration *decreased* 95±174 ms from baseline (*t*(81) = −4.96, *p* < 0.001, *d* = −0.54), with no effects of *session type* or *session order* (*p*’s > 0.34; Figure 4A). Speakers’ peak intensity, maximum pitch, and pitch range did not change from baseline (*p*’s > 0.12), also with no effects of *session type* or *session order* (*p*’s > 0.21; **Figures S3-S5**). To explore how these effects unfolded over two experimental sessions, we re-normalized these metrics to participants’ first-session baselines (Figure 4B), paralleling the exploratory analysis of vowel spacing. Here, sentence duration showed a *session type* × *session order* interaction (*F*(1,81) = 4.73, *p* = 0.04), but no post-hoc pairwise comparisons were significant (*p*’s > 0.08) and adapt-first and control-first participants ended their second sessions with nearly identical durational losses. With increasing familiarity, speakers continued to speed up throughout their second session: adapt-first participants were 106±175 and then 176±246 ms faster; control-first participants were 125±189 and then 178±289 ms faster. The corresponding analyses of intensity and pitch showed, again, no change from baseline (*p*’s > 0.10) and no effects of *session type* or *session order* (*p*’s > 0.24; **Figures S3-S5**). The transfer words showed no changes from baseline in any of the four clear-speech metrics (*p*’s > 0.14).

**Figure 4.**
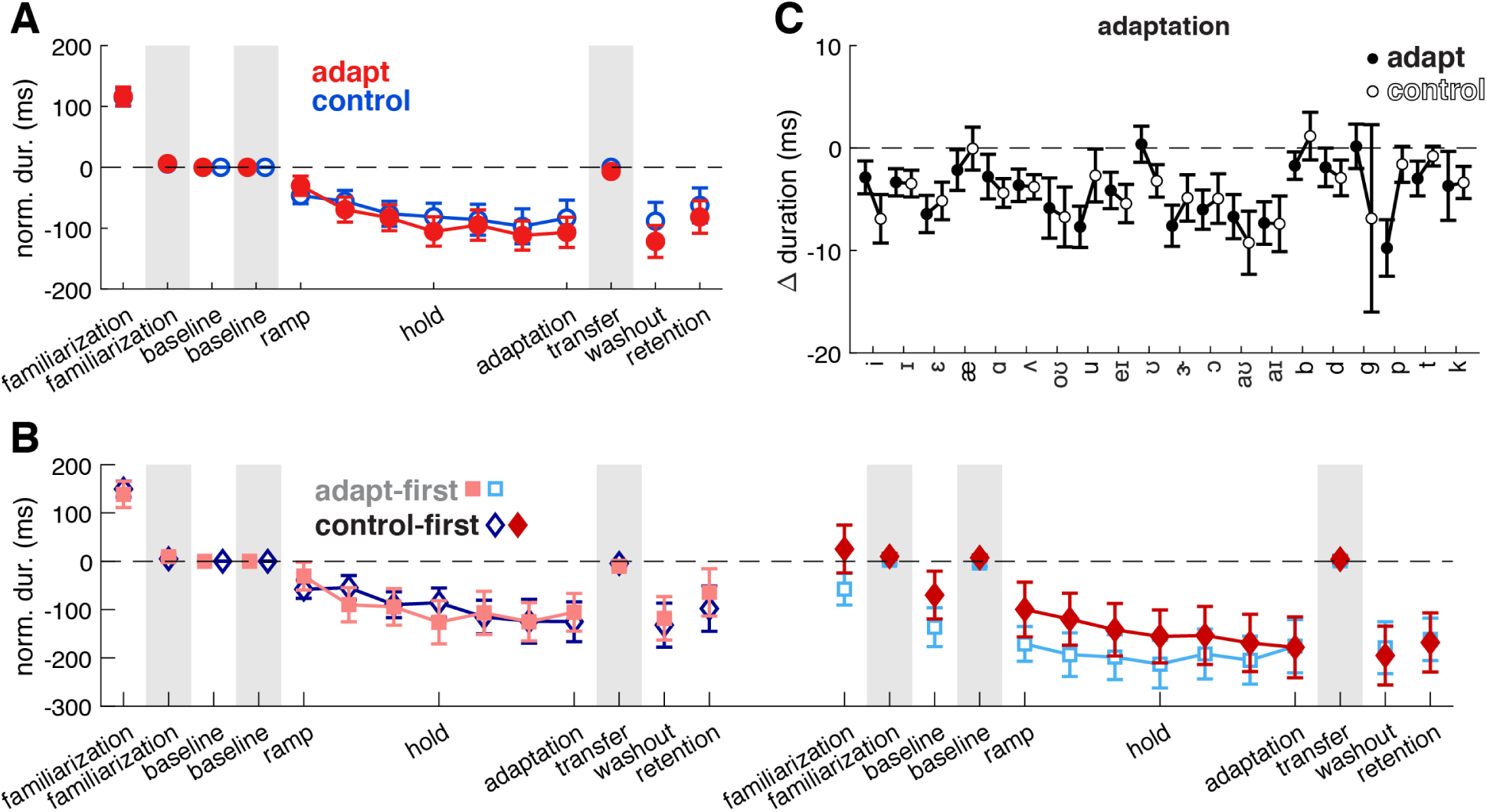
Sentence durations decrease over time. **(A)** Duration collapsed across first and second sessions and normalized to the within-session baseline shows that sentence duration, but not transfer-word duration (shaded regions), decreases in both adapt (red) and control (blue) sessions. **(B)** Duration for first and second sessions separately, normalized to the first-session baseline, reveals accelerating production of sentences in both adapt and control sessions. **(C)** Duration of individual vowel and consonant segments extracted from sentence productions and normalized to the within-session baseline shows that individual sounds do not increase in duration in either adapt (filled) or control (open) sessions. Markers and error bars as in Figure 2.

If utterances with expanded vowel contrast could be described as clear speech, we would also expect to observe lengthening of individual vowel and stop-consonant (/b/, /d/, /g/, /p/, /t/, /k/) sounds (Smiljanić & Bradlow, 2008; Picheny et al., 1986) in the adapt session relative to the control session. In fact, *no* vowels increased in duration from baseline to the adaptation phase, and 12 of 14 sentence vowels *decreased* in duration (*p*’s < 0.01; Figure 4C), all remaining significant after correction for multiple comparisons. The corresponding ANOVA showed this main effect of *segment* (*F*(13,1147) = 3.40, *p* < 0.001) and, critically, no effects of *session type* or *session order* (*p*’s > 0.48). Moreover, stop-consonant durations also decreased 3±21 ms from baseline (*t*(491) = −3.00, *p* < 0.01, *d* = −0.14; Figure 4C), with no effects of *session type*, *session order*, or *segment* (*p*’s > 0.06).

Overall, decreases in sentence and segment durations over time, in combination with no changes in other acoustic cues to prominence, was not consistent with clear speech, but rather with repetition reduction due to familiarity and/or fatigue, affecting both adapt and control sessions and particularly the second of two sessions. In sum, producing multiple repetitions of the same sentences drove participants to speak more quickly; in the face of this, sensorimotor adaptation – not hyperarticulation – expanded vowel contrast, or at least prevented its contraction.

### Perturbation awareness and strategy use do not drive vowel contrast

As a final check, we evaluated speakers’ awareness of the perturbation and whether they responded to it using any explicit strategies that could have contributed to the observed vowel-spacing effects. When queried after their second session, 28 of 41 participants (68%) thought they had ever received a perturbation (as opposed to veridical feedback) and, of those 28, 19 (68%) correctly identified in which session the perturbation had occurred. Although a number of participants reported perceiving various distortions, only two of 41 participants described the manipulation as affecting vowel sounds (**Table S1**). While 21 of 41 speakers reported developing a strategy during the task (e.g., “memorized the sentences”, “switch up the emphasis”, “count trials”), only three of these individuals described a strategy related to pronunciation and/or enunciation (**Table S2**).

To ensure that these five speakers (two “aware” and three “strategic”) did not drive our effects, we repeated our main analyses without them (**Table S3**). In brief, the main effect of *session type* on AVS held for sentences (*F*(1,215) = 6.15, *p* = 0.02), as well as for transfer words (*F*(1,71) = 9.42, *p* < 0.01). Moreover, the vowel-specific results were the same or stronger in the smaller sample (**Table S4**). Thus, observed changes in vowel spacing were not due to explicit pronunciation strategies.

### Vowel contrast predicts intelligibility; contrast gains predict intelligibility gains

Having confirmed that implicit sensorimotor adaptation effected specific changes to speech production, we evaluated whether enhanced vowel contrast translated to increased intelligibility. We focused the perceptual assays on speech produced during second sessions because of the more dramatic acoustic differences between adapt and control conditions there (Figure 2C). Listeners (n = 164) were recruited online and randomly assigned to transcribe one instance of each of the 40 sentences produced by one of the speakers. To mitigate effects of listener ability, each listener heard, in random order, 20 productions taken from the baseline phase and the other 20 taken from the adaptation phase. To simulate conditions of reduced speech intelligibility and ensure that we could measure change from baseline given our sample population of healthy (and presumably, highly intelligible) speakers, the productions were embedded in speech-shaped noise at −21 dB SNR, verified during pilot testing to result in ∼75% word recognition at baseline. Intelligibility was calculated as the percentage of words transcribed correctly.

At baseline, speakers with greater AVS were more intelligible (*r* = 0.37, *p* = 0.02; Figure 5A), underscoring the relationship between vowel contrast and effective communication. Critically, the change in AVS from baseline to adaptation was positively associated with changes in intelligibility (*r* = 0.36, *p* = 0.02; Figure 5B). Of note, however, many speakers’ intelligibility actually *decreased* as the experiment progressed. We hypothesized that the accelerated production of familiar sentences (i.e., **Figure 4**’s significant decrease in sentence duration) could explain the loss of intelligibility. However, even after entering the *change in duration* into a stepwise linear regression model with speaker *age*, speaker *sex*, and the *change in AVS*, the *change in AVS* remained the only significant predictor of changes in intelligibility (*F*(1,39) = 5.84, *p* = 0.02, R^2^ = 0.13). The model estimated that for every 1% increase in AVS, the speaker could expect 0.44 percentage points of gain in sentence intelligibility.

**Figure 5.**
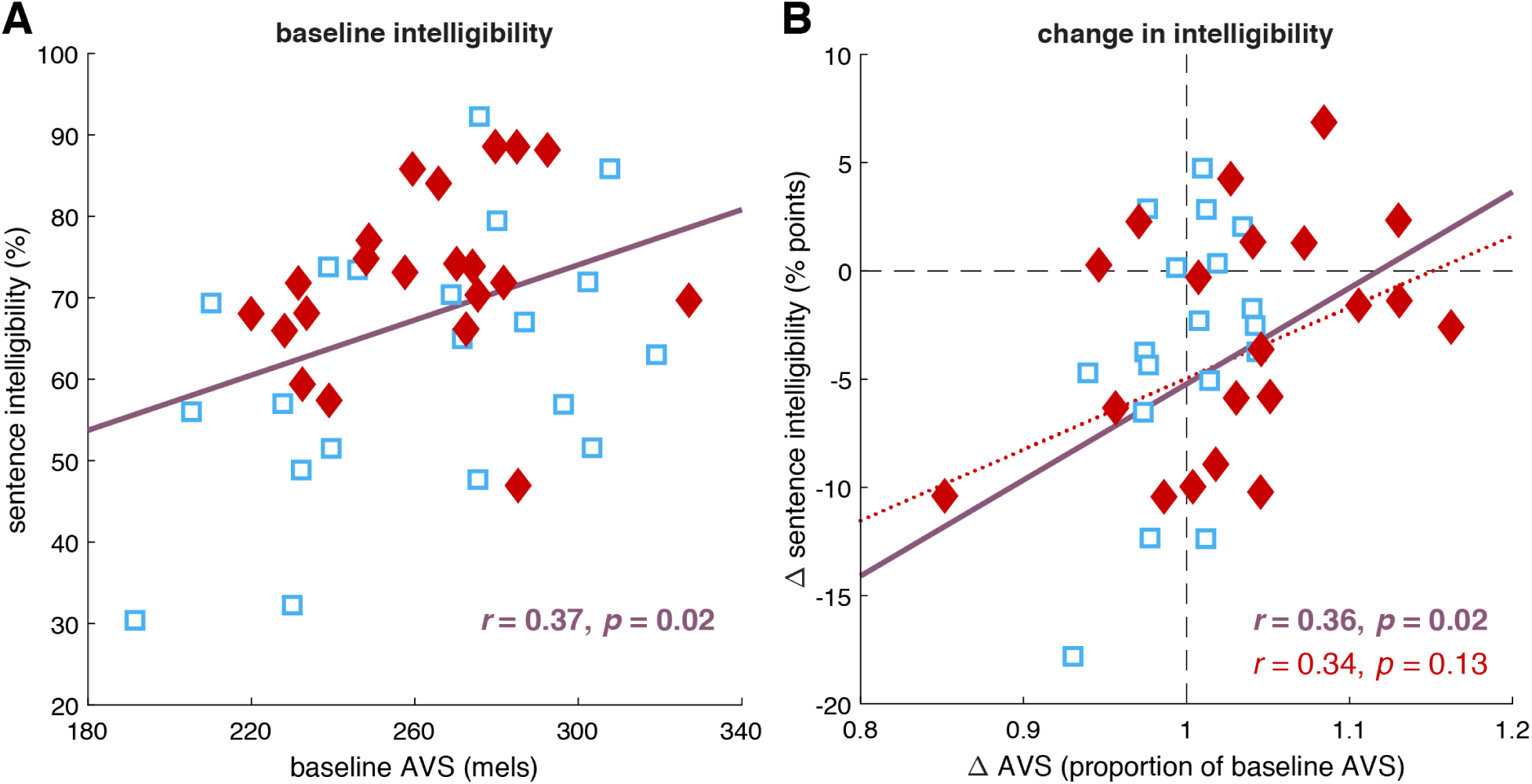
AVS predicts speech intelligibility. **(A)** At baseline, AVS is positively associated with the intelligibility of sentences in noise. **(B)** Gains in AVS from baseline to the adaptation phase are positively associated with gains in sentence intelligibility over the same period. Red diamonds: adapt-session participants. Blue squares: control-session participants. Purple least-square lines and correlations: computed over both participant groups. Red least-squares line and correlation: computed over adapt-session participants. All data are from second sessions.

To understand whether the relationships between vowel spacing and intelligibility were robust to different metrics, we repeated the analyses in Figure 5 using VSA and AAVS instead of AVS. Baseline VSA was not associated with baseline intelligibility, (*r* = 0.25, *p* = 0.11), nor were gains in VSA associated with changes in intelligibility from baseline (*r* = 0.10, *p* = 0.55). Baseline AAVS was positively associated with baseline intelligibility, (*r* = 0.36, *p* = 0.02), but gains in AAVS were not significantly associated with changes in intelligibility from baseline (*r* = 0.18, *p* = 0.25). These results roughly parallel those quantifying adaptation, with AVS and AAVS being more sensitive than VSA to the acoustic and perceptual outcomes of interest. Finally, we verified that the three participants who mentioned pronunciation strategies did not drive the intelligibility gains; without them, gains in AVS were still associated with changes in intelligibility from baseline (*r* = 0.37, *p* = 0.02).

It was also the case that baseline AVS measured from the transfer words predicted their baseline intelligibility (*r* = 0.33, *p* = 0.03). In this assay, listeners heard their assigned speaker’s transfer-word productions embedded in speech-shaped noise at −25 dB SNR (chosen via pilot testing to also achieve ∼75% accuracy at baseline) and indicated which word they heard in a nine-alternative forced-choice task. However, there was no significant relationship between the change in AVS due to transfer and the change in transfer-word intelligibility (*r* = 0.19, *p* = 0.23). Indeed, there was no significant change in transfer-word intelligibility across phases (paired *t*(40) = 0.20, *p* = 0.84, *d* = 0.01), likely because vowels in isolated word production are generally much more contrastive (Keating & Huffman, 1984; Farnetani & Faber, 1992; DiCanio et al., 2015). Together, these findings suggest that the communicative benefits of vowel-space expansion mainly accrue to connected speech.

## Discussion

During speech production, we learn from perceived errors by adjusting our speech to oppose those errors, a process known as *sensorimotor adaptation*. Across domains, sensorimotor adaptation has historically been studied with unidirectional shifts applied to simple, ballistic movements separated into individual trials, whether for speech (isolated words), upper limb control (center-out reaches), or eye movement (saccades). Here, we aimed to align the outcomes of adaptation with changes known to increase speech intelligibility – specifically, adaptations that increase the acoustic contrast between vowels in connected speech. Building on prior work that confirmed the role of auditory error processing in complex, natural utterances (Lametti et al., 2018), we altered the auditory feedback that speakers received during sentence production to push all vowels toward the center of vowel space, making them sound less distinct from one another (Parrell & Niziolek, 2021). Indeed, speakers opposed the perturbation and increased global acoustic vowel contrast, with those gains in contrast uniquely predicting gains in intelligibility over the same time period. These findings demonstrate that sensory predictions are highly specific to parts of the acoustic workspace traversed during continuous movements, and that those continuous movements can be adapted to complex, nonuniform perturbations during high-level ecological behavior. We underscore that, because the direction and magnitude of the feedback shift were dependent on current formant values, speakers had to learn multiple opposing transformations, and that these new vowel-specific movements robustly generalized to new kinematic contexts (e.g., the same vowel in different syllables). Furthermore, these articulatory excursions were achieved without a decrease in rate (cf. Perkell et al., 2002; Picheny et al., 1986; Lam et al., 2012), and indeed in the face of increasingly rapid and less distinct production of familiar sentences. Overall, this study establishes the capacity of sensorimotor adaptation to rapidly and implicitly enhance vowel contrast in connected speech.

Similar increases in vowel contrast are a common target of clinical interventions for motor speech disorders, such as the hypokinetic dysarthria associated with Parkinson’s disease (Darley et al., 1969; Pinto et al., 2004; Enderby, 2013), as they are related to increases in speech intelligibility (Weismer et al., 2001; Lam & Tjaden, 2016; Sapir et al., 2007; Tjaden et al., 2013; Whitfield & Goberman, 2014; see also Kain et al., 2007). Several such interventions explicitly instruct patients to adopt and practice a style of speech that is “clear” (e.g., Park et al., 2016; Shin et al., 2022). Clear vs. habitual speech is characterized by greater vowel contrast (Uchanski, 2005; Smiljanić & Bradlow, 2009), which prompts the question of whether the effects we measured could be due not to implicit adaptation, but to purposeful hyperarticulation. To address this concern, we verified that the handful of participants who reported some awareness of the vowel-centralization perturbation and/or pronunciation strategies did not drive our effects (and indeed their behavior was not consistent with either adaptation or hyperarticulation). Additionally, along with vowel spacing, we measured other acoustic markers of clear speech. Three of these markers (peak vocal intensity, maximum pitch, and pitch range) never changed from baseline. The fourth, sentence duration, significantly decreased across two sessions and under both control and adapt conditions, strongly implying that speech was becoming more fluid and natural rather than more careful and controlled. Furthermore, hyperarticulation is nearly synonymous with *increases* rather than decreases in segment duration, providing time to reach more distant articulatory targets (Uchanski et al., 1996; Lindblom, 1990); however, in our data, *no* vowels or stop consonants, the classes of speech segments most frequently described as lengthening under clear speech (Picheny et al., 1986; Krause & Braida, 2004; Ferguson & Kewley-Port, 2007; Smiljanić & Bradlow, 2008; Lam et al., 2012), were lengthened in the adapt session relative to the control session, and indeed the great majority of them were shortened with respect to baseline in both sessions. Altogether, the evidence was inconsistent with purposeful and/or hyperarticulated clear speech causing increases in vowel contrast.

Rather, the learning in the adapt session showed hallmarks of sensorimotor adaptation: it persisted after the perturbation was removed, indicating an update to motor plans, not merely a reflexive response to altered feedback (Raharjo et al., 2021); it was observed in the presence of masking noise, where such reflexive corrections are not possible; and it generalized beyond words in the perturbation stimulus set, suggesting that adaptation was achieved upstream of specific word-level plans (Caudrelier et al., 2018). Moreover, vowel-specific adaptation, as measured by raw increases in distance from the center of the vowel space, was strongly associated with the raw magnitude of the perturbation applied to each vowel (Daliri et al., 2020; Katseff et al., 2012; MacDonald et al., 2010), consistent with a mechanism underpinning vowel adaptation that detects and corrects sensory prediction error (Daliri & Dittman, 2019; Tourville et al., 2008; Chen et al., 2021; Kim et al., 2023). Because this is an unconscious response, sensorimotor adaptation has the potential to drive behavioral change in patients who can exert limited cognitive (Goldman et al., 2018; Zgaljardic et al., 2003) and/or respiratory (Huber & Darling, 2011; Aquino et al., 2021) effort towards modifying their speech.

It is worth emphasizing that the present study was conducted in healthy speakers. Even in this group, those with greater baseline AVS proved to be more intelligible to unfamiliar listeners, underscoring the importance of vowel contrast to speech intelligibility (Bradlow et al., 1996; Neel, 2008; Kewley-Port et al., 2007; Kim et al., 2011). Additionally, the smaller an individual’s baseline AVS, the more AVS they gained, suggesting that learning capitalized on underutilized articulatory-acoustic space and that speakers who have more room to expand may experience the greatest gains in contrast and intelligibility. Given that adult-onset motor speech disorders progress in concert with general physiological declines in speech-related subsystems, such as oromotor muscle strength (Baum & Bodner, 1983; Robbins et al., 1995) and auditory acuity (Cruickshanks et al., 1998; Gadkaree et al., 2016), we included participants across the lifespan and found no evidence that formant adaptation is limited by healthy aging. At present, it seems that sensorimotor adaptation of speech is relatively preserved across the lifespan as well as (if somewhat diminished) in dysarthria (Mollaei et al., 2013; Mollaei et al., 2016; Polsterer, 2024; see also Tsay et al., 2022 and Bock & Schneider, 2002). Together, this evidence suggests that sensorimotor adaptation to vowel centralization may benefit persons who present with reduced vowel spaces.

Of note for basic and applied speech researchers, the choice of vowel-space metric impacted our results (Thompson et al., 2023; Fox & Jacewicz, 2017; Story & Bunton, 2017; Kim et al., 2011). Consistent with our previous study (Parrell & Niziolek, 2021), AVS was sensitive to adaptation to vowel centralization, as it incorporates all inter-vowel distances. VSA, based on data from only four vowels, was less sensitive to sentence-based adaptation. Finally, AAVS, reflecting overall formant dispersion in speech trajectories, showed the largest gains due to adaptation, a 10% increase from baseline. These metrics also revealed the *loss* of vowel contrast over time under typical (control) conditions, which is to be expected in repetitive experiments. Vowel-space contraction, faster speech, and reports of having memorized the sentences together indicate that participants spoke increasingly naturally throughout our experiment. This is important because it represents somewhat of a middle ground between scripted and spontaneous speech, which is massively acoustically reduced (Johnson, 2004; van Bergum & Koopmans-van Beinum, 1989). Future studies should directly test whether sensorimotor adaptation can drive increases in vowel contrast in spontaneous speech and conversation.

In sum, we have shown that auditory perturbations to vowel distinctiveness can drive speakers to increase vowel contrast in connected speech, with positive effects on their intelligibility. This study is important for models of sensorimotor integration across all motor domains, as it shows that complex, nonuniform alterations to sensory feedback during continuous movement can not only be learned, but successfully applied to drive changes relevant to ecological behavior. The rapid, implicit, and robust nature of this learning suggest it as a promising approach to the motor rehabilitation of developmental and neurologic conditions that impair speech intelligibility (Roemmich & Bastian, 2018).

## Acknowledgements

The authors thank Rob Olson at the Waisman Center’s Clinical Translational Core for technical assistance. This work was supported by NIH grant R01 DC019134 to C.A.N. and B.P.

## Author contributions

C.A.N. and B.P. designed research. S.A.J. performed experiments and processed the data. S.D.B. analyzed the data and wrote the manuscript. S.D.B., C.A.N., and B.P. revised the manuscript. All authors approved the final version.

## Declaration of interests

The authors declare no competing interests.

## Data and code availability

Data have been deposited at https://osf.io/3fhbg/ and are publicly available as of the date of publication. Original code has been deposited at https://github.com/blab-lab/vsaSentence and is publicly available as of the date of publication.

## METHODS

### Participants (speakers)

Forty-two participants across three age bands were recruited for this study: n = 14 in the younger group (18 to 30 years), n = 14 in the middle-aged group (31 to 55 years), and n = 14 in the older group (56 years and older). One participant in the older group was excluded from analysis due to difficulty reading the stimulus words and producing a large number of speech errors. The final sample of n = 41 included 10 males and 31 females ranging in age from 18 to 76 years. Information about gender identity was reported by 24 of the 41 participants; in all cases, it was identical to their report of sex assigned at birth. All participants reported their race as White; one participant additionally reported their race as Black. One participant reported Hispanic ethnicity. As a proxy for socioeconomic status, the educational background of the sample was as follows: high school graduate (n = 3), some college (n = 13), two-year college graduate (n = 7), four-year college graduate (n = 11), and post-graduate (n = 7). All participants were native speakers of American English and reported no history of speech, hearing, or neurological disorders. Auditory thresholds were tested using the modified Hughson-Westlake procedure (Hughson & Westlake, 1944; Carhart & Jerger, 1959). All thresholds in the younger and middle-aged participants were ≤25 dB HL; for the older participants, some hearing loss at 4000 Hz (≤50 dB HL) was allowed. Informed consent was obtained, and participants received either monetary compensation or course credit. The Institutional Review Board of the University of Wisconsin–Madison approved all study procedures.

### Participants (listeners)

One hundred sixty-four participants were recruited via the online research platform Prolific (https://www.prolific.com/) and received monetary compensation. The study was only made available to registered participants who met the following prescreening criteria: aged 18 to 45 years; located in the United States; first and primary language is English; fluent in English; no language- or literacy-related disorders; no diagnoses of attention deficit or autism spectrum disorders; no neurological disorders or conditions; no history of head injury; normal hearing; and normal or corrected to normal vision. The sample included 88 males and 76 females with a mean age of 31 years (SD = 7 years). Information about gender identity was not provided. The racial/ethnic background of the sample was as follows: Asian (n = 25), Black (n = 28), Mixed (n = 14), Other (n = 5), White (n = 91), and no data (n = 1). As a proxy for the socioeconomic status of those who reported their activities, participants were working full time (n = 85), working part time (n = 23), not in paid work (n = 13), unemployed (n = 24), or students (n = 36). Listeners (who remained anonymous) were provided with a consent form and indicated their agreement to participate before being directed to the website where the experiment was hosted. The Institutional Review Board of the University of Wisconsin–Madison approved all study procedures.

### Speech production experiment

#### Auditory perturbation

Speakers’ auditory feedback was recorded, altered, and played back using Audapter (Cai et al., 2008; Tourville et al., 2013; https://github.com/shanqing-cai/audapter_matlab), a software program that performs real-time manipulation of acoustic parameters of speech. Participants’ speech was recorded at 48 kHz with an AKG C520 head-mounted microphone, digitized with a Focusrite Scarlett sound card, and delivered to a desktop computer. Audapter downsampled the signal to 16 kHz, identified the first and second vowel formants (F1 and F2) via linear predictive coding (LPC), and filtered the signal to introduce the desired formant shift. For speech to which no shift was applied, Audapter returned the unmodified signal with the same processing delay (∼18 ms). Participants received their auditory feedback through Beyerdynamic DT 770 closed-back circumaural headphones. Speech was played back at ∼80 dB SPL and mixed with speech-shaped noise at 60 dB SPL that served to mask unaltered speech that would otherwise have been potentially perceptible via air or bone conduction. (The level of speech playback (∼80 dB SPL) is approximate because the microphone gain was set to send a signal of 80 dB SPL based on the participant’s habitual speech volume in the calibration phase; participants were of course free to speak more or less loudly in subsequent trials.) During transfer-word trials, instead of speech, masking noise was played at 78 dB SPL over the headphones to mask any perception of unaltered speech.

The formant shift in this study was dependent on current values of F1 and F2 and was applied continuously during the production of connected speech. A speaker-specific perturbation field was calculated such that all vowels were pushed towards the center of the speaker’s vowel space. This center was defined as the centroid of the quadrilateral formed by the four corner vowels of English (/i/, /æ/, /ɑ/, and /u/) as produced in isolated words during the calibration phase of the experiment (see below). The magnitude of the perturbation was defined as a percentage of the distance between the current formants and the center; this magnitude gradually increased during the ramp phase and was then held at a maximum of 50%.

#### Stimuli and trial structure

Sentence stimuli included 40 phonetically balanced sentences selected from the Harvard sentences (IEEE, 1969; https://www.cs.columbia.edu/~hgs/audio/harvard.html) commonly used in speech research. Transfer words were the following: *bead*, *bid*, *bayed*, *bed*, *bad*, *bod*, *bud*, *hood*, *bode*, and *booed*, selected to cover the majority of the English vowel space. Stimuli were presented one at a time on an LED computer screen. Participants were asked to read the item aloud as it appeared. Each sentence was presented for 4.5 s and each word for 1.5 s. The interstimulus interval was randomly jittered between 0.75 and 1.5 s. Sentence production was organized into 40-trial blocks, with each sentence appearing once per block in random order. Transfer-word presentation was organized into 50-trial blocks composed of five ten-trial groups, with each word appearing once per group in random order.

#### Procedure

This was a two-visit study in which speakers completed a session with (“adapt”) and a session without (“control”) altered auditory feedback. The session order was randomized and counterbalanced; n = 20 participants were “adapt-first” and n = 21 were “control-first.” The purpose of the control session was to understand acoustic changes that may occur over the course of an hour of speaking. All participants completed both sessions, with approximately one week between them (mean = 8.56 days; mode = 7; range = 6 to 33).

The phase structure of each production session was as follows (Figure 1B):

– Calibration: Two blocks of transfer words masked by noise; no perturbation; used to define a participant-specific LPC order for formant tracking in Audapter and to identify the center of the vowel space.
– Familiarization: One block of sentences; no perturbation.
– Familiarization: One block of transfer words masked by noise; no perturbation.
– Baseline: One block of sentences; no perturbation; used as reference values.
– Baseline: One block of transfer words masked by noise; no perturbation; used as reference values.
– Ramp: One block of sentences; the perturbation magnitude increased incrementally from 0 to 50% of the distance to center (maximum perturbation).
– Hold: Five blocks of sentences; maximum perturbation.
– Adaptation: One block of sentences; maximum perturbation; used to measure adapted connected speech at its steady state.
– Transfer: One block of transfer words masked by noise; no perturbation; used to measure transfer of adaptation to isolated words never produced with the perturbation.
– Washout: One block of sentences; no perturbation; used to measure aftereffects in connected speech once the perturbation is removed.
– Retention: One block of sentences after a ten-minute pause during which participants were asked not to speak; no perturbation; used to measure short-term retention of adaptation in connected speech.

The total time per session was ∼65 minutes. A self-paced break was offered every 20 to 25 trials. At the end of the second visit, speakers completed a brief questionnaire that assessed their awareness of the perturbation as well as any strategies they used during the study.

### Speech perception experiment

#### Stimuli and trial structure

In order to evaluate speakers’ baseline intelligibility and change in intelligibility, a subset of their utterances were presented to unfamiliar listeners for tests of speech reception. Four listeners were assigned to each speaker; each listener heard only one speaker. The critical comparison was within-speaker, between utterances produced prior to the onset of the centralization perturbation and those produced after sustained exposure to the perturbation. All stimuli for the perception experiment came from speakers’ second sessions; thus, n = 21 speakers contributed adapt-session data and n = 20 speakers contributed control-session data.

To assess the reception of isolated transfer words differing only in vowel nucleus, listeners identified which word they heard in a nine-alternative forced-choice task. The transfer word “hood” was not used in this task as the initial /h/ would provide a cue to word identity. The remaining 45 items from each of the baseline phase and the transfer phase were used, for a total of ten trials of each of nine words. These items were presented in random order, interspersed with ten catch trials (not analyzed other than to verify listeners’ attention and effort), with no indication that items were taken from different phases or differed from one another in any way.

To assess the reception of connected speech, listeners transcribed sentences. To prevent perceptual learning effects, we established a counterbalancing scheme such that each listener heard each sentence only once. Specifically, for each speaker, we randomly divided the 40 sentences into two groups of 20, A and B. Listeners 1A and 2A heard the A sentences from the baseline phase and the B sentences from the adaptation phase. Listeners 1B and 2B heard the B sentences from the baseline phase and the A sentences from the adaptation phase. The 40 sentences were presented to each listener in random order with no indication that they were taken from different phases or differed from one another in any way.

Sentence and transfer-word productions were excluded from the perception experiment if they contained errors. Errorful productions were flagged by trained research assistants who viewed the visual stimulus while listening to the speech recording for each trial. Transfer-word trials were excluded if the speaker uttered the wrong word or a severe distortion. Sentence trials were excluded if the speaker uttered any wrong word and/or performed an audible self-correction. Short pauses were allowed but other disfluencies were not. In addition, trials were excluded if the recording ended before the participant had finished speaking. After excluding errorful transfer-word productions, all 90 trials remained for n = 37 of the speakers; n = 3 speakers yielded 88 trials and n = 1 speaker yielded 84 trials. After excluding errorful sentence productions, an average of 79 out of a possible 80 items (40 baseline and 40 adaptation) remained for each speaker (SD = 1, range = 75 to 80).

Because the intelligibility of unimpaired adult speakers is near 100%, stimuli were mixed with speech-shaped noise in order to degrade overall intelligibility (Bent et al., 2022; Dubno et al., 2002; Summers & Molis, 2004) and permit measurement of potential perceptual improvements. The signal-to-noise ratio was determined through pilot testing to achieve ∼75% intelligibility across vowels at baseline: for transfer-word stimuli, this was −25 dB SNR, and for sentence stimuli, −21 dB SNR. Catch trials were generated from baseline transfer-word productions (one instance of each word and two of *bud*) and presented at a more favorable −7 dB SNR. To produce the noise-masked stimuli, the audio recording for each correct trial was automatically segmented into its component phonemes by the Montreal Forced Aligner (McAuliffe et al,, 2017; https://montreal-forced-aligner.readthedocs.io), verified by a research assistant who reviewed the waveform, spectrogram, and aligned text grid. For the monosyllabic transfer words, we calculated the intensity of the vowel segment using the root mean square of the signal from vowel onset to vowel offset. We then calculated the intensity of a segment of speech-shaped noise of equal duration in the same way, and scaled the speech signal to achieve the desired SNR using the formula SNR = 20×log(signal/noise). The noise was added to the scaled vowel signal, a buffer of 200 ms before and 200 ms after the vowel was added, and a rising and falling ramp of 10 ms was applied to the final audio file. The sentence stimuli were generated in the same way, except that the speech signal was delimited by the onset of the first phone and the offset of the final phone in the sentence.

#### Procedure

The structure of the perception session (self-paced, ∼25 minutes in length) was as follows:

– <u>Volume calibration 1</u>: To calibrate computer volume to the perceived loudness of the noise bursts used in the headphone checks, listeners were asked to go to a quiet area, connect their headphones, and set their computer volume “a bit below 50%.” They were then instructed to play a noise stimulus sound file and increase their volume until it was “loud but not uncomfortable.” Listeners attested to hearing the sound clearly before they could advance.
– <u>Headphone screening 1</u>: After a guided example trial, listeners completed six trials of the Huggins Pitch test (Milne et al., 2021; https://app.gorilla.sc/openmaterials/100917). This test asks listeners to identify which of three white noise sounds contains a faint tone; the tone percept is induced when the listener is wearing headphones but not when they are listening over speakers.
– <u>Headphone screening 2</u>: After a guided example trial, listeners completed six trials of the Beats test (Milne et al., 2021; https://app.gorilla.sc/openmaterials/100917). This test asks listeners to identify which of three sounds is the smoothest. A percept of beats is generated when the listener is using speakers, but the percept is smooth when listening over headphones.
– Volume calibration 2: To calibrate computer volume to the perceived loudness of the upcoming speech stimuli, listeners were asked to play a sound file containing a familiar two-syllable word embedded in speech-shaped noise at −15 dB SNR, to adjust their volume until it was “loud but not uncomfortable,” and to type the word before they could advance.
– <u>Transfer words</u>: Listeners heard one noise-masked item per trial and clicked a button to indicate which word they heard. Word buttons were present on screen throughout the experiment in a 3 × 3 grid. The mapping of each word to a button was randomized across listeners (e.g., which word occupied the top left button), but the mapping was fixed throughout the experiment for each listener. The task was self-paced, as listeners clicked “Play” to initiate a 500-ms pause followed by the auditory stimulus. Each stimulus was able to be played only once, and a response was required before the next stimulus could be played. A progress bar indicated the proportion of trials completed.
– <u>Sentences</u>: Listeners heard one noise-masked sentence per trial and typed what they heard in a text box. The task was self-paced, as listeners clicked “Play” to initiate a 500-ms pause followed by the auditory stimulus. Each stimulus was able to be played only once, and a response (at minimum, entering a space) was required before the next stimulus could be played. A progress bar indicated the proportion of trials completed.

### Analysis

#### Vowel formants

Formants, or resonances of the vocal tract, provide the primary cues to vowel identity. Formant data were tracked using wave_viewer (Niziolek & Houde, 2015; https://github.com/blab-lab/wave_viewer), a Matlab interface to Praat software (Boersma & Weenink, 2019; http://www.praat.org/). LPC order and pre-emphasis values were set for each participant, as was an amplitude threshold that indicated the onset of vocalization. The Montreal Forced Aligner (McAuliffe et al., 2017; https://montreal-forced-aligner.readthedocs.io) was used to automatically segment the signal into its component phones given the stimulus text. The Aligner used American English acoustic models as well as pronouncing dictionaries (i.e., “english_us_arpa”); as such, the vowels identified were the following: AA, AE, AH, AO, AW, AY, EH, ER, EY, IH, IY, OW, UH, and UW (the vowel OY (/ɔɪ/) did not appear in any of our stimuli). In the manuscript and figures, we referred to these 14 “sentence vowels” using the International Phonetic Alphabet (IPA), respectively: /ɑ/, /æ/, /ʌ/, /ɔ/, /aʊ/, /aɪ/, /ɛ/, /ɝ/, /eɪ/, /ɪ/, /i/, /oʊ/, /ʊ/, and /u/. As expected, the sentence stimuli yielded more tokens for some vowels than for others (*F*(13,12628) = 5812.10, *p* < 0.001), consistent with the overall distribution of phonemes in spoken American English (Lammert et al., 2020). There were no systematic differences in vowel counts in adapt vs. control sessions or by phase of the experiment (all main-effect and interaction *p*’s ≥ 0.73).

The ten transfer vowels (/ɑ/, /æ/, /ʌ/, /ɛ/, /eɪ/, /ɪ/, /i/, /oʊ/, /ʊ/, and /u/) were identified from the production of isolated, monosyllabic transfer words (*bod*, *bad*, *bud*, *bed*, *bayed*, *bid*, *bead*, *bode*, *hood*, and *booed*, respectively) in the same way.

Segmentation was verified by trained research assistants who manually adjusted the vowel boundaries when necessary, referencing the waveform and spectrogram. The mean F1 and F2 values were then extracted from the middle 50% of each vowel and converted from Hz to mels, a perceptually based frequency scale.

#### Vowel-space metrics

Average vowel spacing (AVS) was calculated as the average pairwise distance, in mels, between all 14 sentence vowels and all 10 transfer vowels. Larger AVS indicates greater acoustic contrast among vowels. Vowel space area (VSA) was calculated as the area, in mels^2^, of the quadrilateral formed by the corner vowels /i/, /æ/, /ɑ/, and /u/ in F1-F2 space. Larger VSA indicates a larger working vowel space. Articulatory-acoustic vowel space (AAVS) was calculated as the square root of the generalized variance of all of a speaker’s F1 and F2 values in each phase. Larger AAVS indicates a greater range of articulatory-acoustic motion (Whitfield & Goberman, 2014).

All three vowel-space metrics were normalized to facilitate comparisons across time and session types. The primary normalization in this study was with respect to the participant’s within-session baseline. The mean raw vowel-space metric in each phase of sentence production and transfer-word production was divided by the mean raw baseline value for the corresponding stimulus (sentences or transfer words). Thus, values >1 indicate expansion and values <1 indicate contraction. In an exploratory analysis, we also normalized values with respect to the baseline of each participant’s first (chronological) session in the same manner.

Vowel-space metrics were evaluated with repeated-measures ANOVA with fixed factors of *session type* (adapt, control), *session order* (adapt-first, control-first), and *phase* (adaptation, washout, retention), as well as a random factor of *subject* nested within *session order*. Full models with all main effects and interaction terms were calculated. For testing against baseline, *t*-tests were Holm-Bonferroni corrected for multiple comparisons and used Cohen’s *d* as a measure of effect size.

Adaptation (indexed by first-session normalized AVS in the adaptation phase of the speaker’s adapt session) as a function of speaker characteristics (*age*, *sex*, and *baseline AVS* in the first session) was evaluated with stepwise linear regression using a combination of forward and backward selection starting from a constant model. The criterion for adding a predictor was a *p*-value < 0.05 for the *F*-test of a change in the sum of the squared error; the criterion for removing a predictor was a *p*-value > 0.10 for the *F*-test of a change in the sum of the squared error.

#### Vowel-specific metrics

In addition to global measures such as AVS that take all vowels into account, we also quantified how individual vowels moved with respect to the center of vowel space. The participant-specific center was defined as the centroid of the four corner vowels’ locations in F1-F2 space during the production of isolated words in the calibration phase. The Euclidean distance from this center to the mean location of each vowel in each phase of the experiment served as each vowel’s “distance-to-center.” The baseline value of this distance was taken from the respective baseline phases for sentence vowels and transfer vowels separately. Vowel-specific adaptation was measured as the distance-to-center in the adaptation phase minus the baseline value (for sentence vowels) and as the distance-to-center in the transfer phase minus the baseline value (for transfer vowels).

Vowel-specific adaptation was evaluated with repeated-measures ANOVA with fixed factors of *session type* (adapt, control), *session order* (adapt-first, control-first), and *vowel* (with 14 levels for sentences and 10 levels for transfer words), as well as a random factor of *subject* nested within *session order*. Full models with all main effects and interaction terms were calculated. For comparing vowel-specific adaptation between sessions and against baseline, *t*-tests were Holm-Bonferroni corrected for multiple comparisons and used Cohen’s *d* as a measure of effect size.

The magnitude of vowel-wise adaptation was modeled as adaptation ∼ perturbation + (1+perturbation|subject) + (1+perturbation|vowel), where perturbation was calculated as the mean perturbation size, in mels, over all tokens of the given vowel in the adaptation phase of participants’ adapt session. In this phase of the experiment, the perturbation size was fixed at 50% of the Euclidean distance from current F1 and F2 values to the center of the participant’s vowel space.

#### Clear-speech metrics

For each sentence and transfer-word production, we extracted four acoustic parameters related to clear speech: duration (in ms), peak intensity (calculated from the root mean square Audapter signal and reported in arbitrary units), maximum vocal pitch (in Hz), and pitch range (in Hz). Each metric was normalized by subtracting its mean baseline value from the mean value in each phase of the experiment, such that values >0 indicate increases and values <0 indicate decreases with respect to baseline.

Clear-speech metrics were evaluated with repeated-measures ANOVA with fixed factors of *session type* (adapt, control), *session order* (adapt-first, control-first), and *phase* (baseline, adaptation), as well as a random factor of *subject* nested within *session order*. Full models with all main effects and interaction terms were calculated. All *t*-tests were Holm-Bonferroni corrected for multiple comparisons and used Cohen’s *d* as a measure of effect size.

#### Intelligibility

Intelligibility was first calculated within each listener and then averaged across the four listeners who were randomly assigned to each speaker.

Sentence intelligibility was measured as the percent of words transcribed correctly, computed separately for sentences produced during the baseline and adaptation phases. For each sentence trial, both the text entered by the listener and the sentence recorded by the speaker were converted to a string of phonemes using a pronouncing dictionary. An orthographic word transcription was scored as correct if it was an exact phonemic match for the spoken target. The total number of correctly transcribed words in each phase was divided by the total number of spoken words across all sentences in that phase.

Transfer-word intelligibility was measured as the percentage of nine-alternative forced-choice trials answered correctly, computed separately for transfer words produced during the baseline and transfer phases.

The bivariate relationships between baseline AVS and baseline intelligibility, and between changes in AVS and changes in intelligibility, were computed with Pearson correlation. Additionally, the potential contributions of *change in AVS*, speaker *age*, speaker *sex*, and *change in duration* to changes in intelligibility were evaluated with stepwise linear regression as detailed above.

## Supplementary Information

**Figure S1.**
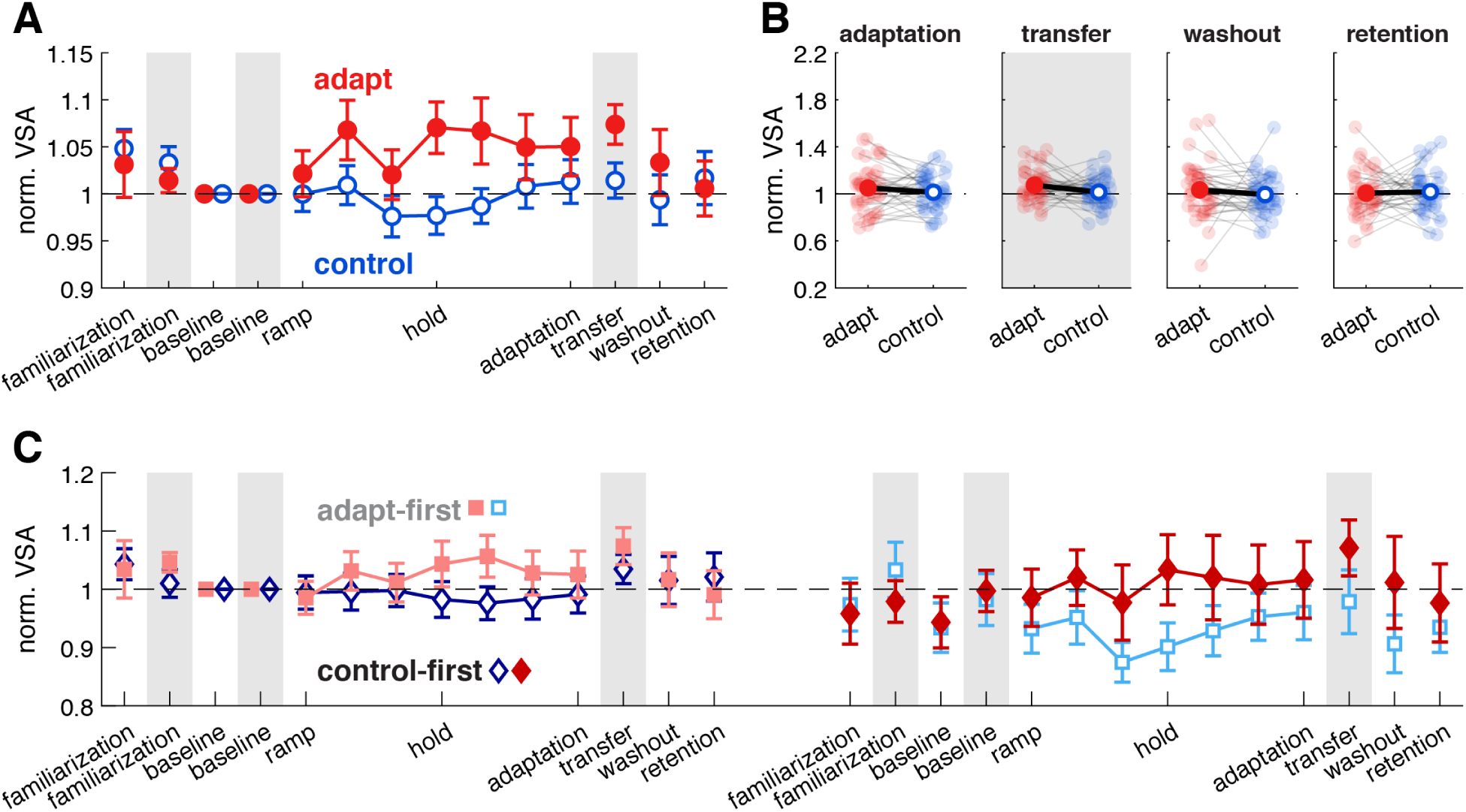
VSA is sensitive to transfer but not to sentence-based adaptation. **(A)** VSA collapsed across first and second sessions and normalized to the within-session baseline does not distinguish adapt (red) vs. control (blue) sentence productions in the adaptation phase, but does indicate generalization of adaptation to untrained transfer words (shaded regions). **(B)** Individual data (small markers connected by gray lines) and group means for within-session normalized VSA in each phase of interest. **(C)** VSA for first and second sessions separately, normalized to the first-session baseline, similarly does not distinguish adapt vs. control in the adaptation phase. As in Figure 2, data from adapt-first participants are pink squares followed by light blue squares. Data from control-first participants are dark blue diamonds followed by red diamonds. Error bars show standard error.

**Figure S2.**
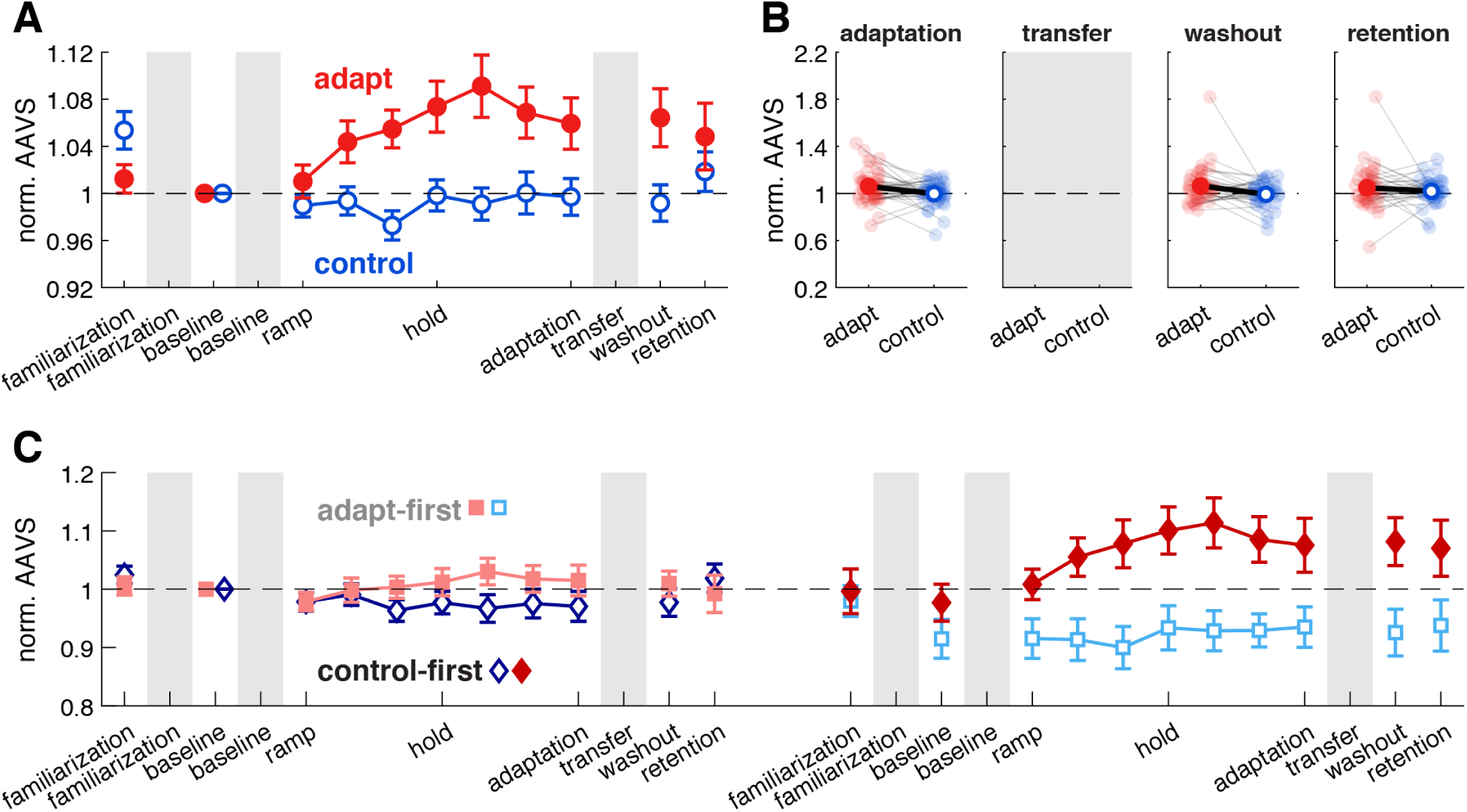
Adaptation increases articulatory working space. **(A)** AAVS collapsed across first and second sessions and normalized to the within-session baseline shows that articulatory working space increases in adapt (red) but not control (blue) sessions. Because AAVS is a trajectory-based measure, it is not well defined for the isolated word productions in the transfer phase (thus no data in shaded regions). **(B)** Individual data (small markers connected by gray lines) and group means for within-session normalized AAVS in each phase of interest. **(C)** AAVS for first and second sessions separately, normalized to the first-session baseline, confirms the adaptation effect. Markers and error bars as in Figures 2 and S1.

**Figure S3.**
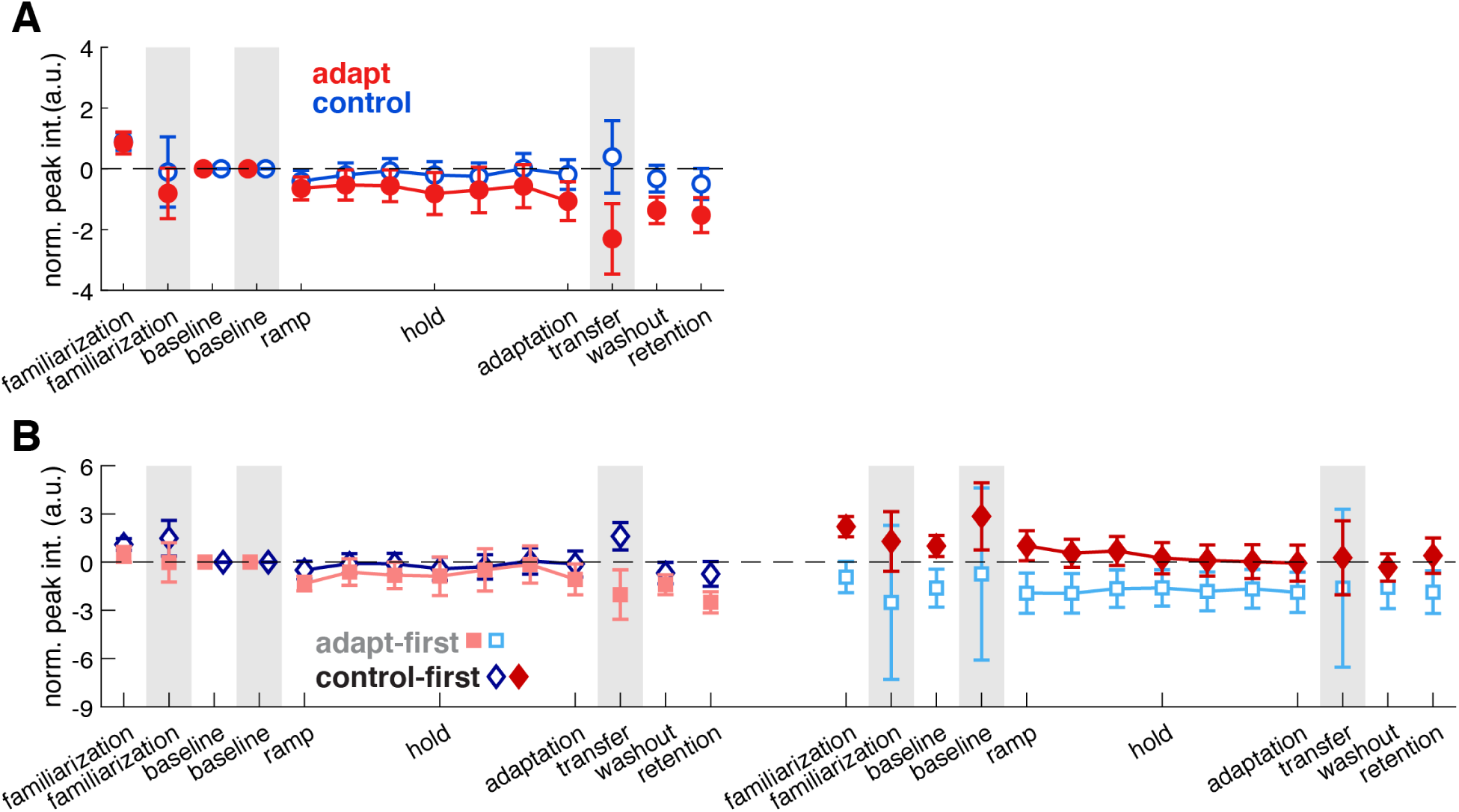
Peak intensity is stable over time. **(A)** Peak intensity, in arbitrary units, collapsed across first and second sessions and normalized to the within-session baseline shows no change from baseline for either sentences or transfer words (shaded regions), in either adapt (red) or control (blue) sessions. **(B)** Peak intensity for first and second sessions separately, normalized to the first-session baseline, reveals no changes over the course of two sessions. Markers and error bars as in Figures 4 and S1.

**Figure S4.**
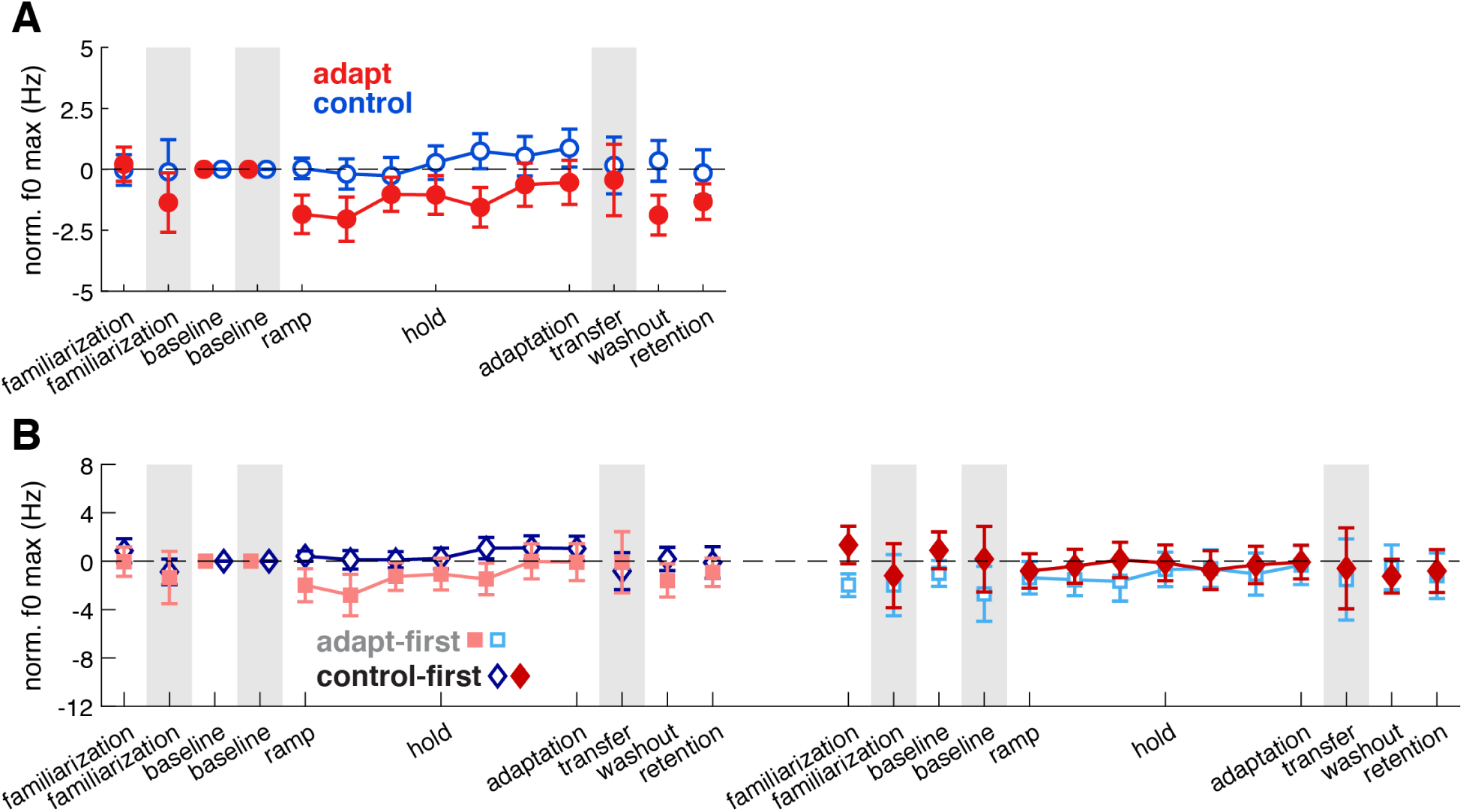
Maximum vocal pitch (f0) is stable over time. **(A)** Maximum pitch collapsed across first and second sessions and normalized to the within-session baseline shows no change from baseline for either sentences or transfer words (shaded regions), in either adapt (red) or control (blue) sessions. **(B)** Maximum pitch for first and second sessions separately, normalized to the first-session baseline, reveals no changes over the course of two sessions. Markers and error bars as in Figures 4 and S1.

**Figure S5.**
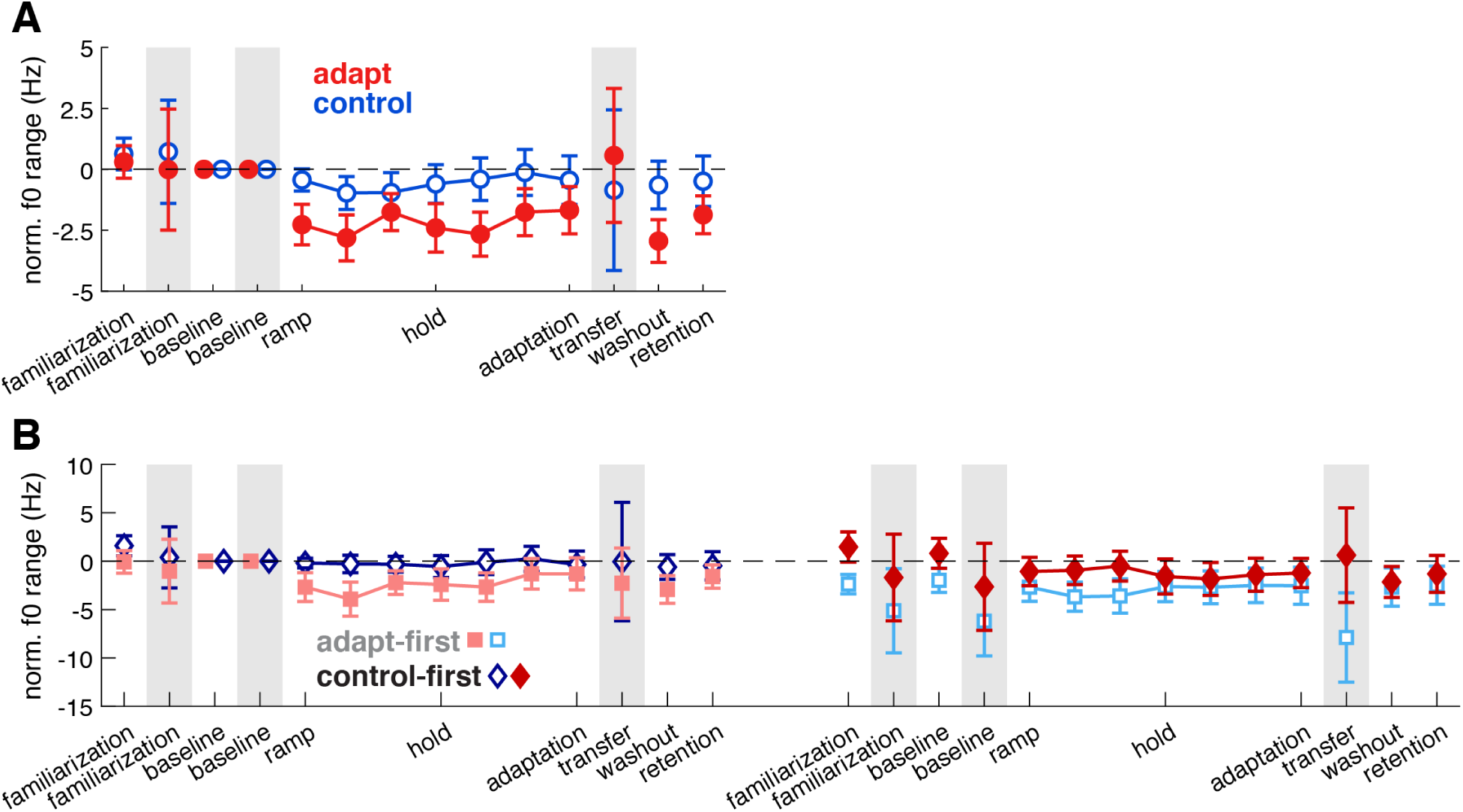
Vocal pitch (f0) range is stable over time. **(A)** Pitch range collapsed across first and second sessions and normalized to the within-session baseline shows no change from baseline for either sentences or transfer words (shaded regions), in either adapt (red) or control (blue) sessions. **(B)** Pitch range for first and second sessions separately, normalized to the first-session baseline, reveals no changes over the course of two sessions. Markers and error bars as in Figures 4 and S1.

**Table S1.**
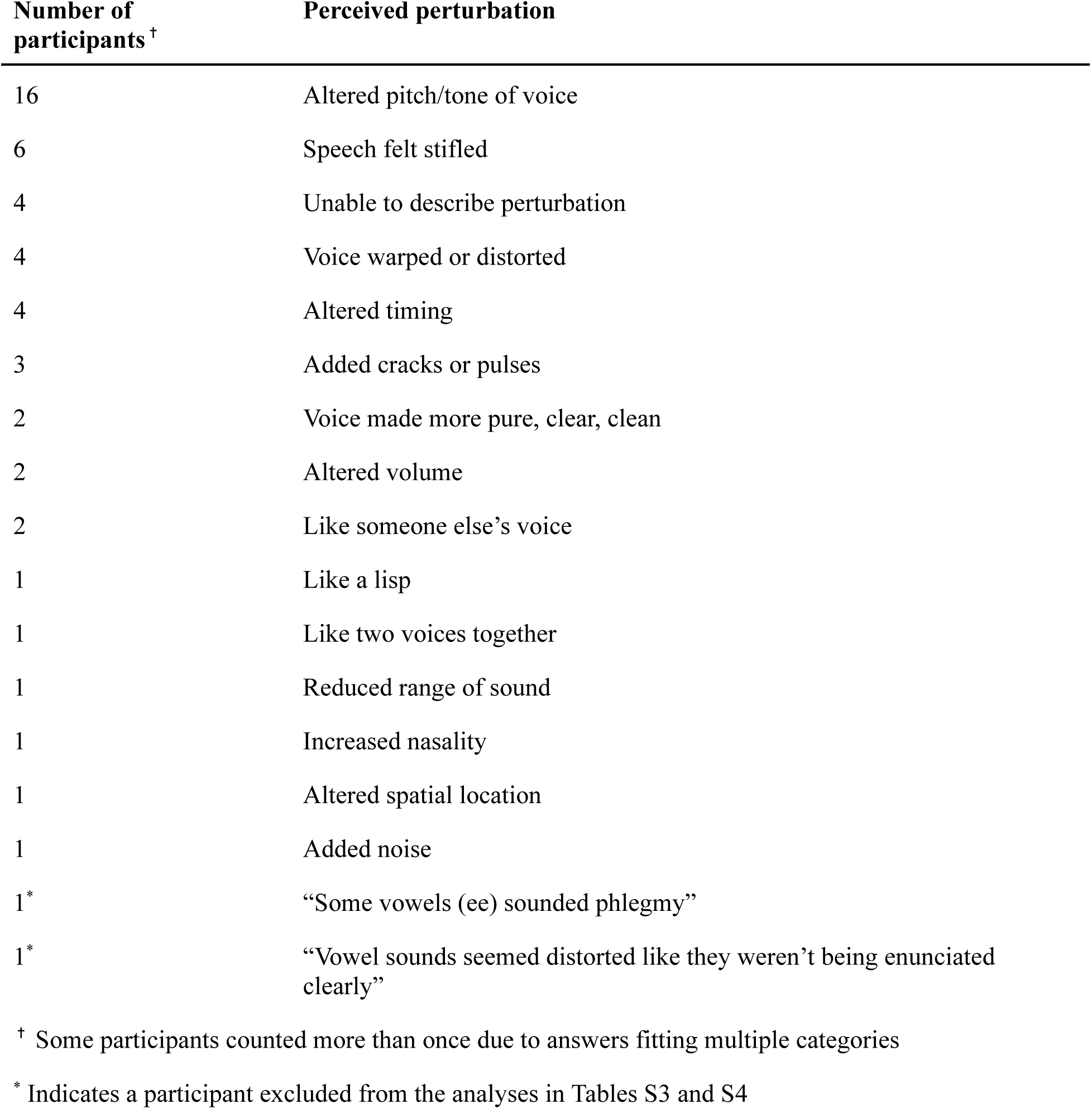
Perturbation awareness reported by participants.

| **Number of participants<sup>†</sup>** | **Perceived perturbation** |
| --- | --- |
| 16 | Altered pitch/tone of voice |
| 6 | Speech felt stifled |
| 4 | Unable to describe perturbation |
| 4 | Voice warped or distorted |
| 4 | Altered timing |
| 3 | Added cracks or pulses |
| 2 | Voice made more pure, clear, clean |
| 2 | Altered volume |
| 2 | Like someone else's voice |
| 1 | Like a lisp |
| 1 | Like two voices together |
| 1 | Reduced range of sound |
| 1 | Increased nasality |
| 1 | Altered spatial location |
| 1 | Added noise |
| 1* | "Some vowels (ee) sounded phlegmy" |
| 1* | "Vowel sounds seemed distorted like they weren't being enunciated clearly" |

<sup>†</sup> Some participants counted more than once due to answers fitting multiple categories
\* Indicates a participant excluded from the analyses in Tables S3 and S4

**Table S2.** Strategy use reported by participants.

| **Number of participants** | **Reported strategy** |
| --- | --- |
| 20 | None |
| 4 | Memorized sentences |
| 4 | Varied word emphasis or voice inflection |
| 2 | Stayed focused and awake |
| 2 | Maintained consistency |
| 2 | Ignored audio |
| 2 | Took a break or got up to stretch |
| 1 | Spoke louder |
| 1 | Counted trials on fingers |
| 1* | “The less you hear yourself, the more you enunciate” |
| 1* | “To pronounce as clearly as possible” |
| 1* | “I tried to make the tense vowels particularly clear” |

\* Indicates a participant excluded from the analyses in Tables S3 and S4

**Table S3.** AVS results excluding participants with vowel-perturbation awareness or pronunciation strategies.

| **Variable** | **AVS<sup>†</sup> in Sentences** | **AVS<sup>†</sup> in Transfer Words** |
| --- | --- | --- |
| *session type* | $F(1,215) = 6.15, p = 0.02$ | $F(1,71) = 9.42, p < 0.01$ |
| *phase* | $F(2,215) = 0.98, p = 0.38$ | - |
| *session order* | $F(1,215) = 0.78, p = 0.38$ | $F(1,71) = 0.39, p = 0.54$ |
| *session type* × *phase* | $F(2,215) = 3.05, p = 0.054$ | - |
| *session type* × *session order* | $F(1,215) = 4.15, p = 0.049$ | $F(1,71) = 0.29, p = 0.60$ |
| *phase* × *session order* | $F(2,215) = 1.55, p = 0.22$ | - |
| *session type* × *phase* × *session order* | $F(2,215) = 0.19, p = 0.82$ | - |

<sup>†</sup> AVS normed within session to parallel the analyses in Figure 2A

**Table S4.** Vowel-specific results excluding participants with vowel-perturbation awareness or pronunciation strategies.

| **Change in distance from center<sup>†</sup>** | **Sentences** | **Transfer Words** |
| --- | --- | --- |
| differed between adapt and control | /i/, /ɪ/, /e/, /ʊ/, /aʊ/ | /i/, /ɪ/, /e/, /ɑ/, /ʌ/ |
| differed from baseline | /i/, /ɪ/, /e/, /æ/, /ʌ/, /ʊ/ | /i/, /ɪ/, /e/ |

<sup>†</sup> *t*-tests corrected for multiple comparisons

## Notes

https://osf.io/3fhbg/

### Competing Interest Statement

The authors have declared no competing interest.

### Summary of Updates

New analyses depicted in Figure 4C.

## References

Aquino, Y. C., Cabral, L. M., Miranda, N. C., Naccarato, M. C., Falquetto, B., Moreira, T. S., & Takakura, A. C. (2022). Respiratory disorders of Parkinson’s disease. Journal of Neurophysiology, 127(1), 1–15.

Baker, R. E., & Bradlow, A. R. (2009). Variability in word duration as a function of probability, speech style, and prosody. Language and Speech, 52(4), 391–413.

Baum, B. J., & Bodner, L. (1983). Aging and oral motor function: Evidence for altered performance among older persons. Journal of Dental Research, 62(1), 2–6.

Bent, T., Baese-Berk, M., Ryherd, E., & Perry, S. (2022). Intelligibility of medically related sentences in quiet, speech-shaped noise, and hospital noise. The Journal of the Acoustical Society of America, 151(5), 3496–3508.

Bock, O., & Schneider, S. (2002). Sensorimotor adaptation in young and elderly humans. Neuroscience & Biobehavioral Reviews, 26(7), 761–767.

Boersma, P., & Weenink, D. (2019). Praat: Doing phonetics by computer. (Version 6.0.47) [Computer software]. http://www.praat.org/

Bradlow, A. R., Torretta, G. M., & Pisoni, D. B. (1996). Intelligibility of normal speech I: Global and fine-grained acoustic-phonetic talker characteristics. Speech Communication, 20(3-4), 255–272.

Burnham, D. K., Joeffry, S., & Rice, L. (2010). Computer-and human-directed speech before and after correction. In Proceedings of the 13th Australasian International Conference on Speech Science and Technology, 14-16 December 2010, Melbourne, Australia (pp. 13–17).

Cai, S., Boucek, M., Ghosh, S., Guenther, F. H., & Perkell, J. (2008). A system for online dynamic perturbation of formant trajectories and results from perturbations of the Mandarin triphthong /iau/. Proceedings of the 8th International Seminar on Speech Production, 65–68.

Carhart, R., & Jerger, J. F. (1959). Preferred method for clinical determination of pure-tone thresholds. Journal of Speech and Hearing Disorders, 24(4), 330–345.

Caudrelier, T., Schwartz, J. L., Perrier, P., Gerber, S., & Rochet-Capellan, A. (2018). Transfer of learning: What does it tell us about speech production units?. Journal of Speech, Language, and Hearing Research, 61(7), 1613–1625.

Caudrelier, T., & Rochet-Capellan, A. (2019). Changes in speech production in response to formant perturbations: An overview of two decades of research. Speech Production and Perception: Learning and Memory, 6, 15–75.

Chen, T., Lammert, A. C., & Parrell, B. (2021). Modeling sensorimotor adaptation in speech through alterations to forward and inverse models. In Interspeech (pp. 3201–3205).

Cruickshanks, K. J., Wiley, T. L., Tweed, T. S., Klein, B. E., Klein, R., Mares-Perlman, J. A., & Nondahl, D. M. (1998). Prevalence of hearing loss in older adults in Beaver Dam, Wisconsin: The epidemiology of hearing loss study. American Journal of Epidemiology, 148(9), 879–886.

Daliri, A., & Dittman, J. (2019). Successful auditory motor adaptation requires task-relevant auditory errors. Journal of Neurophysiology, 122(2), 552–562.

Daliri, A., Chao, S. C., & Fitzgerald, L. C. (2020). Compensatory responses to formant perturbations proportionally decrease as perturbations increase. Journal of Speech, Language, and Hearing Research, 63(10), 3392–3407.

Darley, F. L., Aronson, A. E., & Brown, J. R. (1969). Differential diagnostic patterns of dysarthria. Journal of Speech and Hearing Research, 12(2), 246–269.

DiCanio, C., Nam, H., Amith, J. D., García, R. C., & Whalen, D. H. (2015). Vowel variability in elicited versus spontaneous speech: Evidence from Mixtec. Journal of Phonetics, 48, 45–59.

Dubno, J. R., Horwitz, A. R., & Ahlstrom, J. B. (2002). Benefit of modulated maskers for speech recognition by younger and older adults with normal hearing. The Journal of the Acoustical Society of America, 111(6), 2897–2907.

Enderby, P. (2013). Disorders of communication: dysarthria. Handbook of Clinical Neurology, 110, 273–281.

Farnetani, E., & Faber, A. (1992). Tongue-jaw coordination in vowel production: Isolated words versus connected speech. Speech Communication, 11(4-5), 401–410.

Ferguson, S. H., & Kewley-Port, D. (2007). Talker differences in clear and conversational speech: Acoustic characteristics of vowels. Journal of Speech, Language, and Hearing Research, 50(5), 1241.

Fox, R. A., & Jacewicz, E. (2017). Reconceptualizing the vowel space in analyzing regional dialect variation and sound change in American English. The Journal of the Acoustical Society of America, 142(1), 444–459.

Gadkaree, S. K., Sun, D. Q., Li, C., Lin, F. R., Ferrucci, L., Simonsick, E. M., & Agrawal, Y. (2016). Does sensory function decline independently or concomitantly with age? Data from the Baltimore longitudinal study of aging. Journal of Aging Research, 2016(1), 1865038.

Goldman, J. G., Vernaleo, B. A., Camicioli, R., Dahodwala, N., Dobkin, R. D., Ellis, T., … & Simmonds, D. (2018). Cognitive impairment in Parkinson’s disease: a report from a multidisciplinary symposium on unmet needs and future directions to maintain cognitive health. npj Parkinson’s Disease, 4(1), 19.

Hantzsch, L., Parrell, B., & Niziolek, C. A. (2022). A single exposure to altered auditory feedback causes observable sensorimotor adaptation in speech. Elife, 11, e73694.

Houde, J. F., & Jordan, M. I. (1998). Sensorimotor adaptation in speech production. Science, 279(5354), 1213–1216.

Huber, J. E., & Darling, M. (2011). Effect of Parkinson’s disease on the production of structured and unstructured speaking tasks: respiratory physiologic and linguistic considerations. Journal of Speech, Language, and Hearing Research, 54(1), 33–46.

Hughson, W., & Westlake, H. (1944). Manual for program outline for rehabilitation of aural casualties both military and civilian. Transactions of the American Academy of Ophthalmology and Otolaryngology, 48(Suppl), 1–15.

IEEE Recommended Practice for Speech Quality Measurements (1969). IEEE Transactions on Audio and Electroacoustics, 17, 227–246.

Jacobs, C. L., Yiu, L. K., Watson, D. G., & Dell, G. S. (2015). Why are repeated words produced with reduced durations? Evidence from inner speech and homophone production. Journal of Memory and Language, 84, 37–48.

Johnson, K. (2004). Massive reduction in conversational American English. In Spontaneous speech: Data and analysis. Proceedings of the 1st Session of the 10th International Symposium, p. 29–54.

Kain, A. B., Hosom, J. P., Niu, X., Van Santen, J. P., Fried-Oken, M., & Staehely, J. (2007). Improving the intelligibility of dysarthric speech. Speech Communication, 49(9), 743–759.

Katseff, S., Houde, J., & Johnson, K. (2012). Partial compensation for altered auditory feedback: A tradeoff with somatosensory feedback?. Language and Speech, 55(2), 295–308.

Keating, P. A., & Huffman, M. K. (1984). Vowel variation in Japanese. Phonetica, 41(4), 191–207.

Kewley-Port, D., Burkle, T. Z., & Lee, J. H. (2007). Contribution of consonant versus vowel information to sentence intelligibility for young normal-hearing and elderly hearing-impaired listeners. The Journal of the Acoustical Society of America, 122(4), 2365–2375.

Kim, H., Hasegawa-Johnson, M., & Perlman, A. (2011). Vowel contrast and speech intelligibility in dysarthria. Folia Phoniatrica et Logopaedica, 63(4), 187–194.

Kim, K. S., Hinkley, L. B., Dale, C. L., Nagarajan, S. S., & Houde, J. F. (2023). Neurophysiological evidence of sensory prediction errors driving speech sensorimotor adaptation. bioRxiv.

Krause, J. C., & Braida, L. D. (2004). Acoustic properties of naturally produced clear speech at normal speaking rates. The Journal of the Acoustical Society of America, 115(1), 362–378.

Lane, H., Matthies, M., Perkell, J., Vick, J., & Zandipour, M. (2001). The effects of changes in hearing status in cochlear implant users on the acoustic vowel space and CV coarticulation. Journal of Speech, Language, and Hearing Research, 44, 552–563.

Lam, T. Q., & Watson, D. G. (2010). Repetition is easy: Why repeated referents have reduced prominence. Memory & Cognition, 38(8), 1137–1146.

Lam, J., Tjaden, K., & Wilding, G. (2012). Acoustics of clear speech: Effect of instruction. Journal of Speech, Language, and Hearing Research, 55, 1807–1821.

Lam, J., & Tjaden, K. (2016). Clear speech variants: An acoustic study in Parkinson’s disease. Journal of Speech, Language, and Hearing Research, 59(4), 631–646.

Lammert, A. C., Melot, J., Sturim, D. E., Hannon, D. J., DeLaura, R., Williamson, J. R., Ciccarelli, G., & Quatieri, T. F. (2020). Analysis of phonetic balance in standard English passages. Journal of Speech, Language, and Hearing Research, 63(4), 917–930.

Lametti, D. R., Smith, H. J., Watkins, K. E., & Shiller, D. M. (2018). Robust sensorimotor learning during variable sentence-level speech. Current Biology, 28(19), 3106–3113.e2.

Lee, J., Littlejohn, M. A., & Simmons, Z. (2017). Acoustic and tongue kinematic vowel space in speakers with and without dysarthria. International Journal of Speech-Language Pathology, 19(2), 195–204.

Lindblom, B. (1990). Explaining phonetic variation: A sketch of the H&H theory. In: Hardcastle, W.J., Marchal, A. (eds) Speech Production and Speech Modelling. NATO ASI Series, vol 55. Springer, Dordrecht.

Maas, E., Robin, D. A., Hula, S. N. A., Freedman, S. E., Wulf, G., Ballard, K. J., & Schmidt, R. A. (2008). Principles of motor learning in treatment of motor speech disorders. American Journal of Speech-Language Pathology, 17, 277–298.

MacDonald, E. N., Goldberg, R., & Munhall, K. G. (2010). Compensations in response to real-time formant perturbations of different magnitudes. The Journal of the Acoustical Society of America, 127(2), 1059–1068.

McAuliffe, M., Socolof, M., Mihuc, S., Wagner, M., & Sonderegger, M. (2017). Montreal forced aligner: Trainable text-speech alignment using kaldi. Interspeech, 2017, pp. 498–502.

Milne, A. E., Bianco, R., Poole, K. C., Zhao, S., Oxenham, A. J., Billig, A. J., & Chait, M. (2021). An online headphone screening test based on dichotic pitch. Behavior Research Methods, 53, 1551–1562.

Mollaei, F., Shiller, D. M., & Gracco, V. L. (2013). Sensorimotor adaptation of speech in Parkinson’s disease. Movement Disorders, 28(12), 1668–1674.

Mollaei, F., Shiller, D. M., Baum, S. R., & Gracco, V. L. (2016). Sensorimotor control of vocal pitch and formant frequencies in Parkinson’s disease. Brain Research, 1646, 269–277.

Munhall, K. G., MacDonald, E. N., Byrne, S. K., & Johnsrude, I. (2009). Talkers alter vowel production in response to real-time formant perturbation even when instructed not to compensate. The Journal of the Acoustical Society of America, 125(1), 384–390.

Neel, A. T. (2008). Vowel space characteristics and vowel identification accuracy. Journal of Speech, Language, and Hearing Research, 51, 574–585.

Ng, M. L., & Woo, H. K. (2021). Effect of total laryngectomy on vowel production: An acoustic study of vowels produced by alaryngeal speakers of Cantonese. International Journal of Speech-Language Pathology, 23(6), 652–661.

Niziolek, C. A., & Houde, J. (2015). Wave_Viewer: First release. 10.5281/ZENODO.13839.

Oviatt, S., MacEachern, M., & Levow, G. A. (1998). Predicting hyperarticulate speech during human-computer error resolution. Speech Communication, 24(2), 87–110.

Park, S., Theodoros, D., Finch, E., & Cardell, E. (2016). Be Clear: A new intensive speech treatment for adults with nonprogressive dysarthria. American Journal of Speech-Language Pathology, 25(1), 97–110.

Parrell, B., & Niziolek, C. A. (2021). Increased speech contrast induced by sensorimotor adaptation to a nonuniform auditory perturbation. Journal of Neurophysiology, 125(2), 638–647.

Perkell, J. S., Zandipour, M., Matthies, M. L., & Lane, H. (2002). Economy of effort in different speaking conditions. I. A preliminary study of intersubject differences and modeling issues. The Journal of the Acoustical Society of America, 112(4), 1627–1641.

Picheny, M. A., Durlach, N. I., & Braida, L. D. (1986). Speaking clearly for the hard of hearing II: Acoustic characteristics of clear and conversational speech. Journal of Speech, Language, and Hearing Research, 29(4), 434–446.

Pinto, S., Ozsancak, C., Tripoliti, E., Thobois, S., Limousin-Dowsey, P., & Auzou, P. (2004). Treatments for dysarthria in Parkinson’s disease. The Lancet Neurology, 3(9), 547–556.

Polsterer, K. M. (2024). Characterising auditory-motor adaptation of vowel production across age. https://aburlab.web.rug.nl/wp-content/uploads/2024/01/KPolsterer_ThesisArchive.pdf

Purcell, D. W., & Munhall, K. G. (2006). Adaptive control of vowel formant frequency: Evidence from real-time formant manipulation. The Journal of the Acoustical Society of America, 120(2), 966–977.

Raharjo, I., Kothare, H., Nagarajan, S. S., & Houde, J. F. (2021). Speech compensation responses and sensorimotor adaptation to formant feedback perturbations. The Journal of the Acoustical Society of America, 149(2), 1147–1161.

Robbins, J., Levine, R., Wood, J., Roecker, E. B., & Luschei, E. Age effects on lingual pressure generation as a risk factor for dysphagia. Journals of Gerontology Series A: Biological Sciences and Medical Sciences, 50(5), M257–M262.

Rochet-Capellan, A., & Ostry, D. J. (2011). Simultaneous acquisition of multiple auditory–motor transformations in speech. Journal of Neuroscience, 31(7), 2657–2662.

Roemmich, R. T., & Bastian, A. J. (2018). Closing the loop: from motor neuroscience to neurorehabilitation. Annual Review of Neuroscience, 41, 415–429.

Sapir, S., Spielman, J. L., Ramig, L. O., Story, B. H., & Fox, C. (2007). Effects of intensive voice treatment (the Lee Silverman Voice Treatment [LSVT]) on vowel articulation in dysarthric individuals with idiopathic Parkinson disease: Acoustic and perceptual findings. Journal of Speech, Language, and Hearing Research, 50(4), 899.

Sapir, S., Ramig, L. O., Spielman, J. L., & Fox, C. (2010). Formant centralization ratio: A proposal for a new acoustic measure of dysarthric speech. Journal of Speech, Language, and Hearing Research, 53(1), 114–125.

Scarborough, R. (2010). Lexical and contextual predictability: Confluent effects on the production of vowels. Laboratory Phonology, 10, 557–586.

Shin, H., Shivabasappa, P., & Koul, R. (2022). Effect of clear speech intervention program on speech intelligibility in persons with idiopathic Parkinson’s disease: A pilot study. International Journal of Speech-Language Pathology, 24(1), 33–41.

Skodda, S., Visser, W., & Schlegel, U. (2011). Vowel articulation in Parkinson’s disease. Journal of Voice, 25(4), 467–472.

Smiljanić, R., & Bradlow, A. R. (2009). Speaking and hearing clearly: Talker and listener factors in speaking style changes. Language and Linguistics Compass, 3(1), 236–264.

Smiljanić, R., & Bradlow, A. R. (2008). Stability of temporal contrasts across speaking styles in English and Croatian. Journal of Phonetics, 36(1), 91–113.

Story, B. H., & Bunton, K. (2017). Vowel space density as an indicator of speech performance. The Journal of the Acoustical Society of America, 141(5), EL458–EL464.

Summers, V., & Molis, M. R. (2004). Speech recognition in fluctuating and continuous maskers: effects of hearing loss and presentation level. Journal of Speech, Language, and Hearing Research, 47(2), 245–257.

Thompson, A., Hirsch, M. E., Lansford, K. L., & Kim, Y. (2023). Vowel acoustics as predictors of speech intelligibility in dysarthria. Journal of Speech, Language, and Hearing Research, 66(8S), 3100–3114.

Tjaden, K., Lam, J., & Wilding, G. (2013). Vowel acoustics in Parkinson’s disease and multiple sclerosis: comparison of clear, loud, and slow speaking conditions. Journal of Speech, Language, and Hearing Research, 56(5), 1485–1502.

Tourville, J. A., Reilly, K. J., & Guenther, F. H. (2008). Neural mechanisms underlying auditory feedback control of speech. Neuroimage, 39(3), 1429–1443.

Tourville, J. A., Cai, S., & Guenther, F. (2013). Exploring auditory-motor interactions in normal and disordered speech. Proceedings of Meetings on Acoustics, 19(1).

Tsay, J. S., Najafi, T., Schuck, L., Wang, T., & Ivry, R. B. (2022). Implicit sensorimotor adaptation is preserved in Parkinson’s disease. Brain Communications, 4(6), fcac303.

Uchanski, R. M., Choi, S. S., Braida, L. D., Reed, C. M., & Durlach, N. I. (1996). Speaking clearly for the hard of hearing IV: Further studies of the role of speaking rate. Journal of Speech, Language, and Hearing Research, 39(3), 494–509.

Uchanski, R. M. (2005). Clear Speech. The Handbook of Speech Perception, 207–235.

van Bergem, D. R., & Koopmans-van Beinum, F. J. (1989). Vowel reduction in natural speech. In First European Conference on Speech Communication and Technology.

Villacorta, V. M., Perkell, J. S., & Guenther, F. H. (2007). Sensorimotor adaptation to feedback perturbations of vowel acoustics and its relation to perception. The Journal of the Acoustical Society of America, 122(4), 2306–2319.

Weismer, G., Jeng, J. Y., Laures, J. S., Kent, R. D., & Kent, J. F. (2001). Acoustic and intelligibility characteristics of sentence production in neurogenic speech disorders. Folia Phoniatrica et Logopaedica, 53(1), 1–18.

Whitfield, J. A., & Goberman, A. M. (2014). Articulatory–acoustic vowel space: Application to clear speech in individuals with Parkinson’s disease. Journal of Communication Disorders, 51, 19–28.

Zeng, Y., Niziolek, C. A., & Parrell, B. (2023). Simultaneous acquisition of multiple auditory-motor transformations reveals supra-syllabic motor planning in speech production. PsyArXiv.

Zgaljardic, D. J., Borod, J. C., Foldi, N. S., & Mattis, P. (2003). A review of the cognitive and behavioral sequelae of Parkinson’s disease: relationship to frontostriatal circuitry. Cognitive and Behavioral Neurology, 16(4), 193–210.

